# First full-genome alignment representative for the genus *Pestivirus*

**DOI:** 10.1101/2025.05.22.655560

**Authors:** Sandra Triebel, Tom Eulenfeld, Nancy Ontiveros-Palacios, Blake Sweeney, Norbert Tautz, Manja Marz

## Abstract

The members of the genus *Pestivirus* in the family *Flaviviridae* comprise economically important pathogens of life stock like classical swine fever (CSFV) and bovine viral diarrhea virus (BVDV). Intense research over the last years revealed that at least 11 recognized and eight proposed pestivirus species exist. The single-stranded, positive-sense RNA genome encodes for one large polyprotein which is processed by viral and cell-derived proteases into 12 mature proteins. Besides its protein-coding function, the RNA genome also contains RNA secondary structures with critical importance for various stages of the viral life cycle. Some of those RNA secondary structures, like the internal ribosome entry site (IRES) and a 3’ stem-loop essential for genome replication, had already been studied for a few individual pestiviruses.

In this study, we provide the first genome-wide multiple sequence alignment (MSA) including all known pestivirus species (accepted and tentative). Moreover, we performed a comprehensive analysis of RNA secondary structures phylogenetically conserved across the complete genus. While showing well-described structures, like a 5’ stem-loop structure, the IRES element, and the 3’ stem loop SL I to be conserved between all pestiviruses, other RNA secondary structures in the 3’ untranslated region (UTR) were only conserved in subsets of the species. We identified 29 novel phylogenetically conserved RNA secondary structures in the protein-coding region, with so far unresolved functional importance. The microRNA binding site for miR-17 was previously known in species A, B, and C; in this study, we identified it in ten additional species, but not in species K, S, Q, and R. Another interesting finding is the identification of a putative long-distance RNA interaction between the IRES and the 3’ end of the genome. These results together with the now available comprehensive multiple sequence alignment including all 19 pestivirus species, represent a valuable resource for future research and diagnostic purposes.

## Introduction

The members of the genus *Pestivirus* being most closely related to *Hepaci-* and *Pegivirus* within the family *Flaviviridae*, alongside *Orthoflavivirus*, are enveloped, positive-strand RNA viruses with genome lengths ranging from approximately 11.6 to 12.3 kb [1, 2]. Currently, the ICTV lists 11 pestivirus species and 8 more were proposed [3] (see Tab. 1). Pestiviruses like the bovine viral diarrhea viruses (BVDV-1 and −2) and classical swine fever virus (CSFV) are long-known animal pathogens with high relevance to stock farming worldwide [1]. Formerly, pestiviruses were thought to be restricted to cloven-hoofed animals as their hosts. However, when major improvements in bulk sequencing uncovered the sequences of rat pestivirus, Phocoena pestivirus (PhoPeV) from harbour porpoise and pangolin pestivirus as well as the atypical pestiviruses from bats, it became evident that members of this genus infect a much broader host spectrum [3]. The atypical pestivirus from swine (APPV), two novel rodent pestiviruses, and the bat pestiviruses form one monophyletic clade [3]. They differ significantly from the classical pestiviruses listed above (see below). Recently, even more distantly related pestiviruses were identified by protein structure prediction and comparison combined with RNA sequence analysis [2].

The pestiviral genome is organized into a single open reading frame (ORF) flanked by two untranslated regions (UTRs) at the 5’ and 3’ ends. The ORF encodes the polyprotein NH_2_-N^pro^-C-E^rns^-E1-E2-p7-NS2-NS3-NS4A-NS4B-NS5A-NS5B-COO-, with exception of PhoPeV which does not encode for the N-terminal protease N^pro^ [4]. The polyprotein is processed by ER-resident cellular protease as well as the viral proteases N^pro^, non-structural protein 2 (NS2) and NS3/4A into 12 proteins [1]. N^pro^ and the envelope glycoprotein ribonuclease secreted (E^rns^) are only found in pestiviruses and are crucial for combating the host’s innate immune response [5, 6]. The cleavage of NS2-3 into NS2 and NS3 by the cysteine protease in NS2 is vital for pestiviral RNA replication and depends on NS2 protease activation by the cellular chaperone DNAJC14 (formerly termed Jiv) [7]. For NS2 protease activation the Jiv90 domain of DNAJC14 is sufficient [7]. Only the viruses from the APPV-clade, which differ significantly from the classical pestiviruses, especially in their glycoprotein genes (E^rns^, E1 and E2) and the NS2-coding region [8, 9], show no dependency on DNAJC14 [10]. Another peculiarity of pestiviruses is the generation of virus mutants by RNA recombination during persistent infection of ruminants. The cell-derived sequences in the viral genomes encode, e.g., DNAJC14, ubiquitin (Ub), SUMO, LC3, or NEDD3, and are causative for a change in the viral biotype and the induction of a lethal disease in the persistently infected host [11, 12].

**Table 1:**
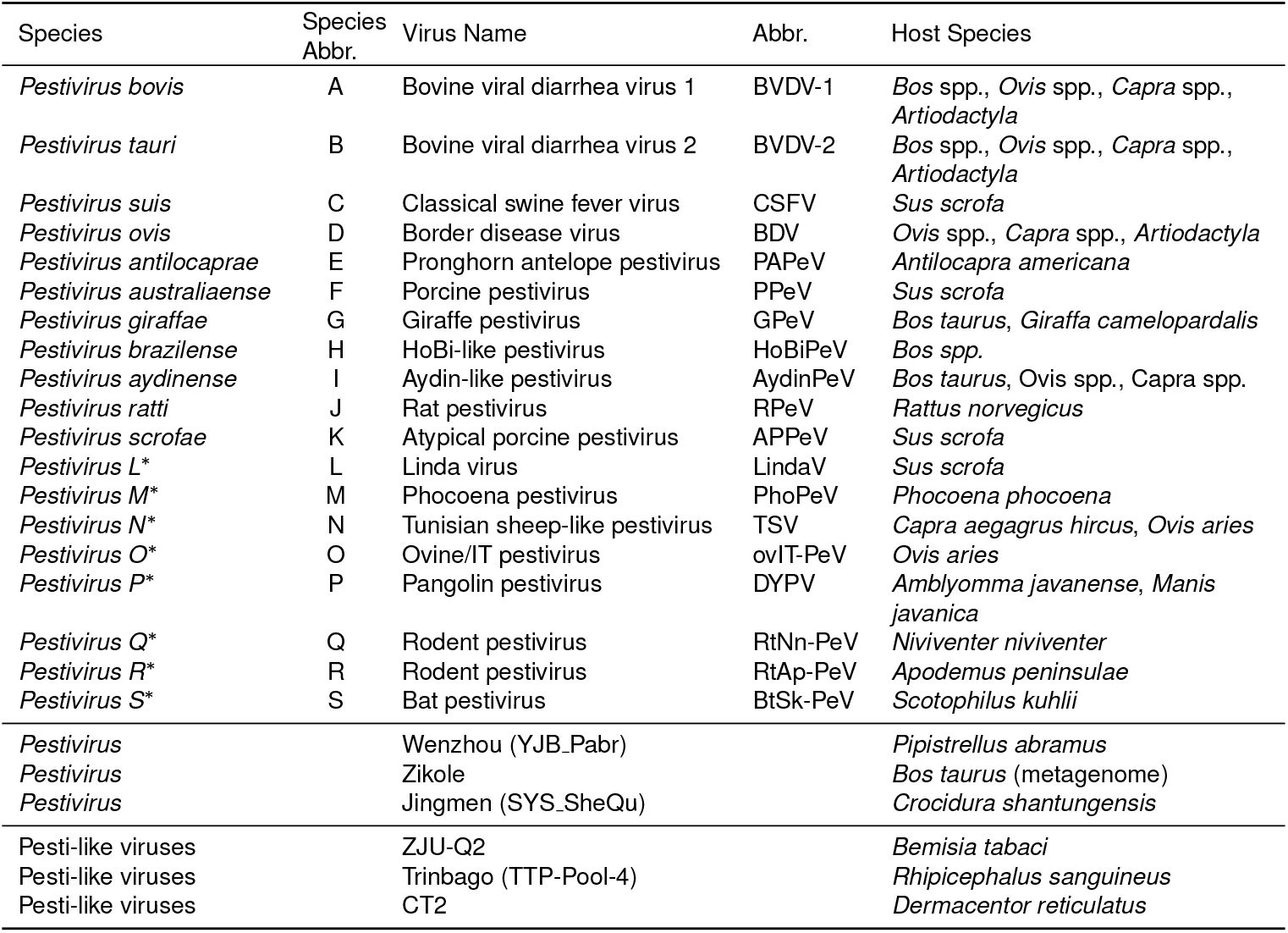
Overview of *Pestivirus* species and corresponding host species. 11 species are already known (A – K), 8 new species (L – S) highlighted with ‘*’ are proposed by Postel *et al*. [3]. Three entries are not assigned to any species but were reported as pestiviruses in our dataset. Three genomes were reported as Pesti-like viruses.

The pestivirus genome neither carries a 5’ cap nor a 3’ poly A tail. Initiation of translation of the viral RNA genome is mediated by a 5’ internal ribosome entry site (IRES), a complex RNA structure recruiting the ribosomes directly to the start codon of the single large open reading frame (ORF) [13, 14]. The pestiviral IRES belongs to type IV analogous to the one in the hepatitis C virus (HCV) genome [15]. Further RNA structures at the 5’ and 3’ ends are crucial for recruiting the viral replicase to the RNA for genome amplification. In the 3’ non-coding region, three stem-loop (SL) structures have been described with the 3’-terminal one, SL I, being of the highest functional relevance [16–18]. More recently, upstream of SL I highly conserved let-7 and miR17 binding sites were identified [19]; miR17 binding was shown essential for replication of BVDV-1 [20].

Here we present the first full-genome alignment of pestiviruses coupled with RNA secondary structure annotation. The alignment was generated using a semi-automated approach and curated by experts in the field of the genus *Pestivirus* and its associated structural elements. In addition to established structures such as the 5’ UTR IRES and 3’ UTR SL I, our study reveals previously unrecognized RNA secondary structures, especially in the protein-coding region, which were predicted by computational methods and thus, are conserved in all pestivirus genomes. The results of our alignment – using genomes covering the entire phylogenetic sequence space of pestivirus isolates – suggest that our predictions are likely to cover virtually all possible RNA secondary structures conserved for all known pestiviruses.

## Material & Methods

### Data

We downloaded a total of 1,684 *Pestivirus* genomes (October 11, 2023) from the `BV-BRC` and `NCBI` databases [21, 22]. To ensure the quality of the data, we filtered the genome status ‘complete’ and excluded the host group ‘lab’.

After identifying redundant genomes utilizing the preprocessing step of our `ViralClust` [23] pipeline, the data set was refined to a total of 756 genomes. Both the original data set and the pre-filtered data set are included in the supplementary files F1 and F2 in `.fasta` format.

### Selection of representative genomes

We clustered the pre-filtered data set based on k-mers to select sequences representing the entire data set: First, we calculated the k-mer profiles of each input sequence. Then we performed a dimension reduction by principal component analysis (PCA), implemented in `scikit-learn` (v1.2.2) [24], followed by clustering using `HDBSCAN` v0.8.27 [25]. We calculated centroid sequences based on the average pairwise distance between all vectors in the same cluster. The sequence with the minimum average distance is selected as the ‘representative genome’ for this cluster. The resulting 52 representative genomes are well distributed throughout the known *Pestivirus* genomes. For comparison, we additionally clustered sequences with these four algorithms: `cd-hit-est` [26, 27], `MMSeqs2` [28], `sumaclust` [29], and `vclust` [30]. The workflow is implemented in `ViralClust` [23]. Based on these results, see Tab. S1, we only used `HDBSCAN` for further analysis. We manually edited our set of representative genomes by (a) adding all 23 RefSeq genomes from `NCBI` [22] and `ICTV` [31, 32], see Tab. S2 ‘RefSeqs’, (b) adding 7 genomes representing outliers or subtrees not covered by the clustering results, see Tab. S2 ‘Outlier’, (c) reducing 23 overrepresented genomes from species A, B, C, and K, see Tab. S2 ‘Reduce overrepresentation’, and (d) removed 1 genome (`JQ799141`) due to a truncated protein. The representative genomes and their distribution throughout the pestiviruses are highlighted in Fig. 1. We visualized the non-redundant dataset in a split graph using `MAFFT` [33] and subsequently `SplitsTree` [34, 35]. The phylogenetic tree is stored in supplementary file F3. For the generation of the alignment presented here, we removed the 3 Pesti-like viruses (`ON684360`, `MN025505`, `MW256672`), since the sequences are over 4,000 nt longer (16,232; 16,274; and 16,802 nt) than the average of the other *Pestivirus* sequences (∼ 12,200 nt) and they show a high phylogenetic distance in the splits graph, see Fig. 1. A total of 55 representative genomes covering all 19 species of pestiviruses were selected, see supplementary file F4.

**Figure 1:**
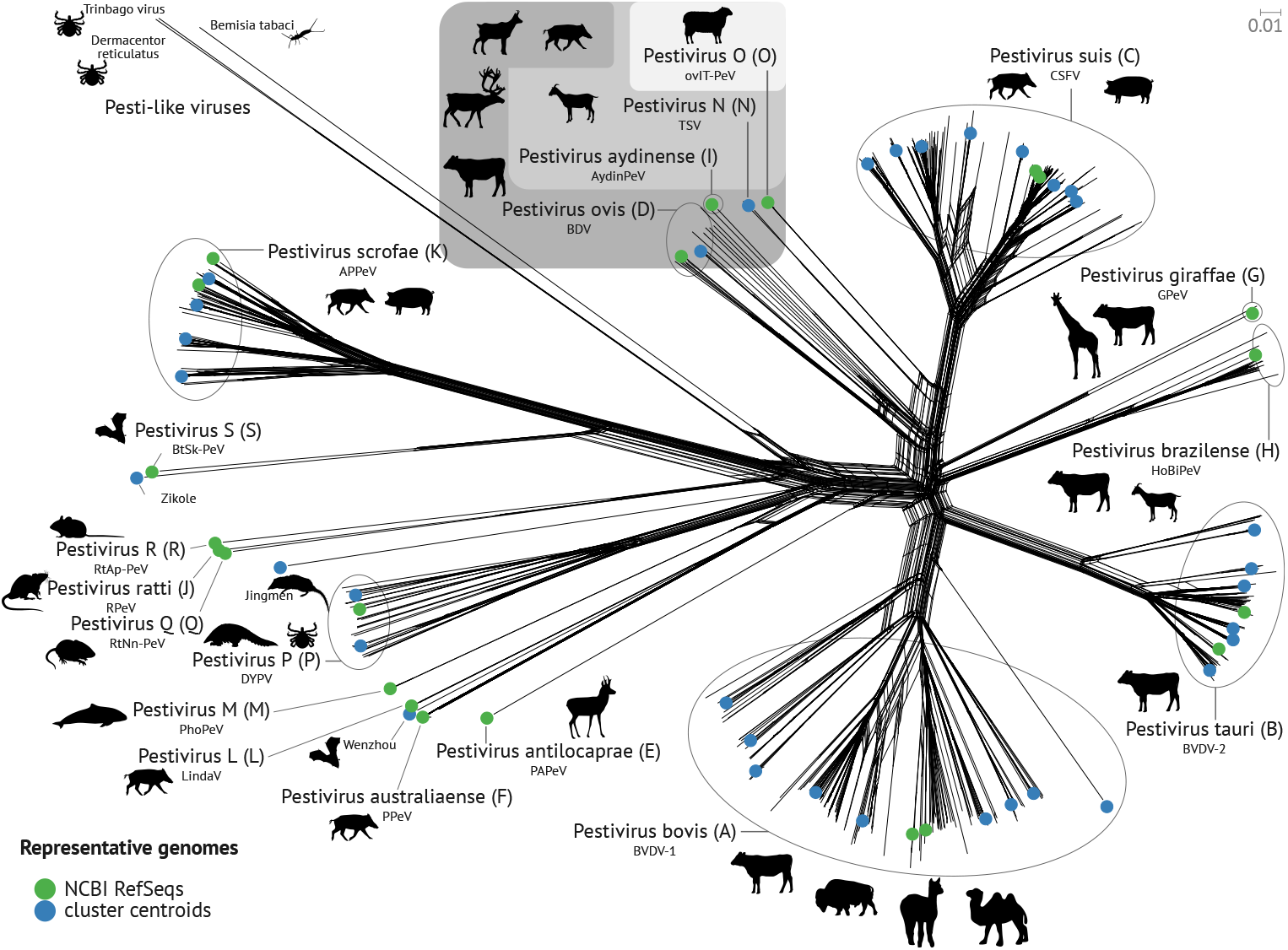
Phylogenetic representation of the *Pestivirus* data set. Our representative genomes consist of NCBI RefSeq genomes (green) and cluster centroids (blue). The tree shows a clear distinction between the different species. The hosts reported for the species are shown by the pictograms. Tab. 1 provides a detailed overview of the hosts. The tree is based on the `MAFFT` alignment of all complete-genome sequences and served as input for `SplitsTree` to construct a splits graph using the neighbor-net algorithm.

**Figure 2:**
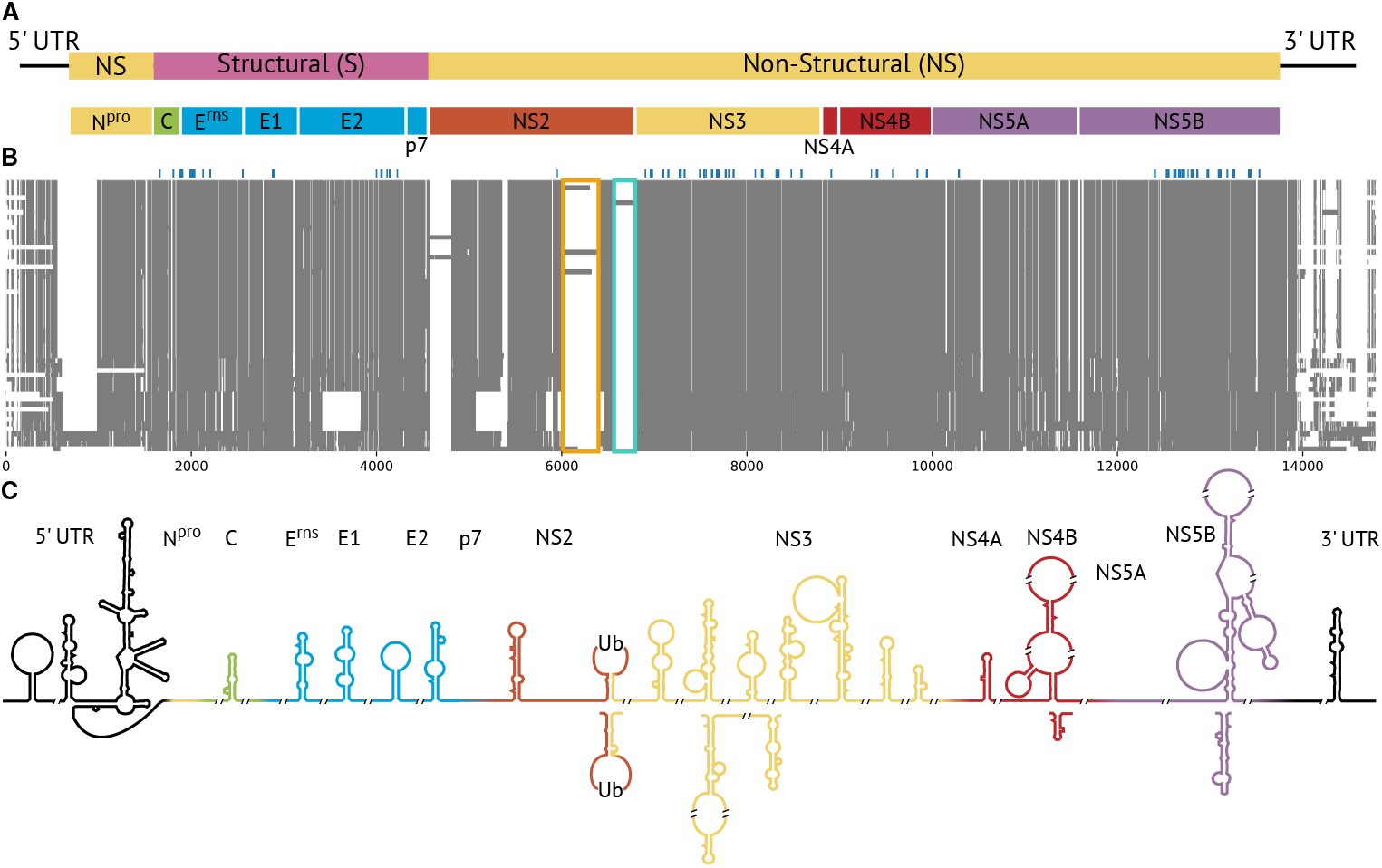
Overview of the (A) genome organization (adapted from ICTV) of *Pestivirus*, (B) full-genome alignment, and (C) RNA secondary structures. Gray areas display the sequences and white areas represent gap regions. Blue above the alignment highlights the location of the anchors calculated by `AnchoRNA`. Colored boxes highlight the insertion of DNAJC14 (orange) and ubiquitin (Ub, cyan).

### Multiple sequence alignment and RNA secondary structure prediction

The final set of 55 representative genomes served as input for constructing the full-genome multiple sequence alignment (MSA), including RNA secondary structure information for the entire sequence. We generated this alignment in five steps by cutting it into sub-alignments, processing, and concatenating them: (1) We identified highly conserved regions (called ‘anchors’) using `AnchoRNA` v1.0.1 [36], see supplementary file F5. Anchors are regions in the translated amino acid sequences present in all genomes, requiring a minimum length of 5, and a minimum BLOSUM62 score of 22 between each anchor in the anchor region and at least one other anchor in the same anchor region. For a detailed description, we refer to Eulenfeld et al. [36]. (2) Subsequently, we focused our analysis on the subregions between these anchors, as well as the 5’ UTR and the 3’ UTR, using `LocARNA` v2.0.0 with additional parameters `--stockholm--consensus-structure alifold` [37]. We temporarily removed for this step 25 highly similar sequences (see Tab. S2 ‘Highly similar sequences’) to bring less bias into the covariance model calculation. (3) The subalignments with the predicted RNA secondary structures were manually curated. At the end of this step, we added the temporarily removed 25 highly similar sequences again to the alignment. (4) We merged the subalignments according to the anchors into one overall MSA. We modified the sequence of `ON165517` in two residues to remove a frameshift in the alignment: (1) changed ‘-’ to ‘N’ in alignment position 3,371; (2) deleted ‘A’ in sequence position 2,672. (5) We searched for RNA secondary structures using a sliding window approach (`RNALalifold`) and analyzed genome circularization using the `ViennaRNA` package [38]. (6) The coding sequence of the nucleotide alignment is then translated into the amino acid alignment.

After we had built the full genome alignment with RNA secondary structure information, we added the annotation of proteins and conserved RNA secondary structures described in the literature. Finally, we selected highly conserved RNA secondary structure regions to export to `Rfam` database [39, 40].

#### Visualization

Alignments were visualized using `Jalview` [41]. In the case of protein alignments, we used the gecos Ocean coloring (based on BLOSUM62) [42]. The overview alignment was visualized using `sugar` [43]. The alignment-based structures were visualized using `R2DT` [44] for the 2D representation and `RNAalifold` [38] for coloring. Single-sequence RNA secondary structure predictions were done using `RNAfold` with parameter `-p` for partition function. If stated, an additional parameter for structural constraints (`-C`) is used [38].

#### Potential generic primer detection

We utilized `AnchoRNA` to find conserved regions on the nucleotide level, which might serve as potential primer in future experiments. Our 55 representative genomes served as input. To be considered a potential primer, we set the parameter for `AnchoRNA` the following: minimum length of 18 nt with a maximum difference of 3 nt to the guiding sequence while at least 50% of the sequences have to contain the conserved region.

## Results & Discussion

In this study, we present a comprehensive analysis of all conserved RNA secondary structures that occur in the complete sequence space of all downloadable pestivirus genomes, see Fig. 1. Thus, the results are supposed to provide a complete description of RNA secondary structures that may be advantageous for the viral life cycle. We found 32 conserved RNA secondary structures present in all pestiviruses, see below. Additionally, we added 6 selected RNA secondary structures available in only subsets of pestiviruses.

For pestiviruses, functionally important RNA secondary structures in the untranslated regions are known. However, with the notable exception of a computational prediction [45], no RNA secondary structures in coding regions have been described so far. A variety of molecular mechanisms can be envisioned in which RNA secondary structure elements are involved. Compared to other *Flaviviridae* RNA secondary structure elements in the protein-coding region, we can imagine the influence on the translation outcome of a particular RNA [46], e.g. by inducing ribosome frameshifts or termination reinitiation. Specific RNA elements can be used to selectively package one RNA, while another, longer RNA species is excluded from packaging by translational inactivation [47]. A translating ribosome may also displace proteins from a secondary structure element of the RNA and thereby initiate a kind of ‘burn after reading’ degradation of the RNA [48]. Therefore, it is important to identify those RNA secondary structures that have been selected for their function from the available sequence space produced by the error-prone replicases of RNA-plus-strand viruses.

### Basic statistics of the full-genome alignments

Our nucleotide-based alignment contains 55 representative *Pestivirus* genomes covering all 19 species (*Pestivirus* A–S). The alignment spans 14,793 residues and 143,147 gaps, averaging approximately 2,603 gaps per sequence. Only four sequences within our alignment contain a total of eleven positions with non-`ACGU` characters (R, M, Y, S, K, and N, see IUPAC code).

The full 5’ UTR exists for 32 sequences (19 incomplete); three sequences lack the 5’ UTR entirely and we manually removed the 5’ UTR sequence of the Jingmen strain (isolate SYS SheQu, accID `OM030319`) due to no sequence or structure similarity. On a structural level, 32 sequences contain domain I (D I), 46 sequences D II, and 48 sequences D III and IV. Domain II, III, and IV form the IRES, see Fig. 3. For the 3’ UTR, 38 sequences display the complete 3’ UTR; three sequences completely lack the 3’ UTR. Only SL I is conserved across all species, see below.

**Figure 3:**
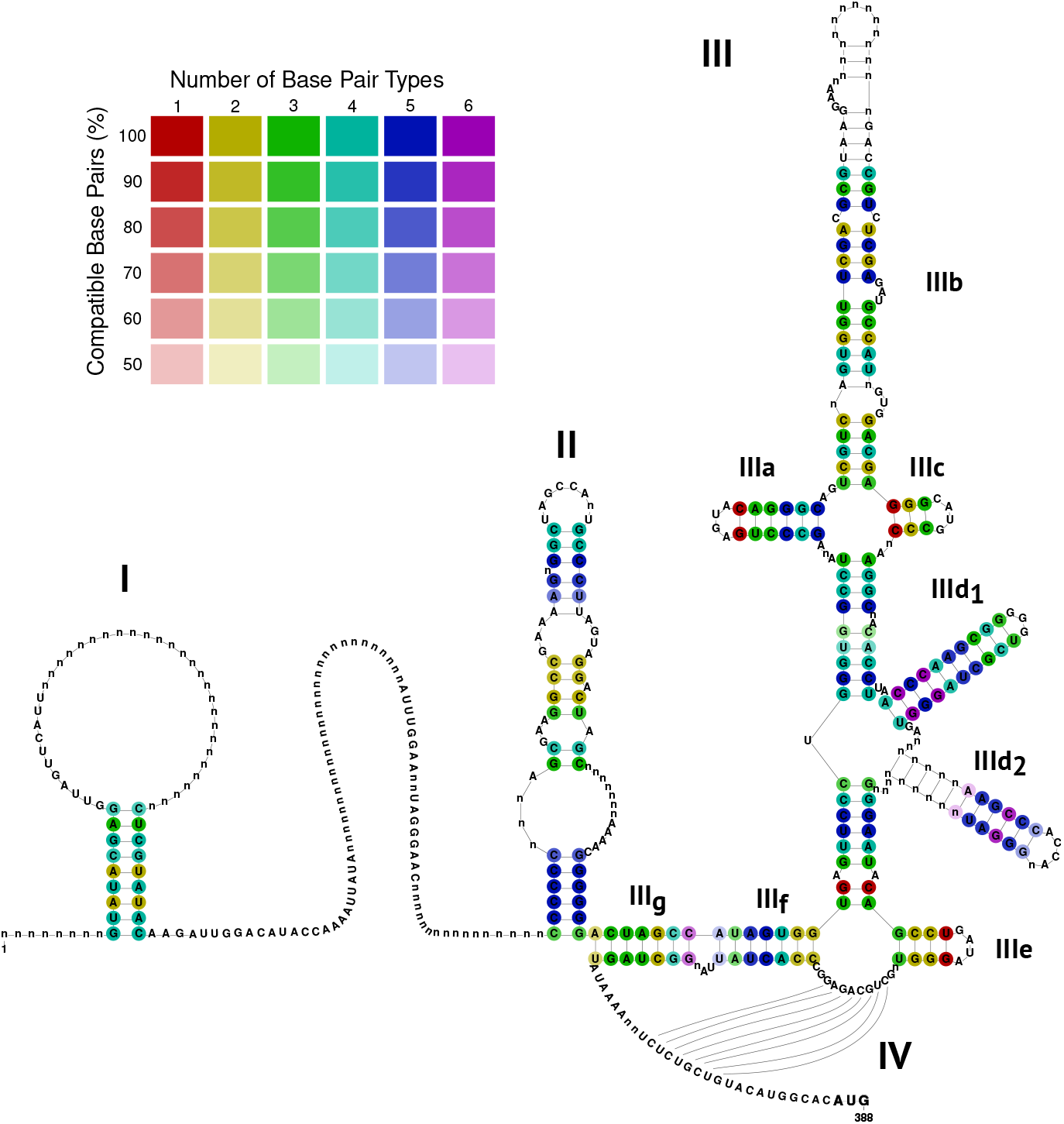
Alignment-based RNA secondary structure prediction of the 5’ UTR, colored by the number of base pair types and the percentage of sequences that form the base pairs, illustrating the extent of covariations in double-stranded regions. Among the selected sequences, 51 isolates had a 5’ UTR sequence (32 complete, 19 incomplete). The 5’ UTR contains D I (present in 32 sequences) and the IRES, which comprises the domains D II (present in 46 sequences), D III, and D IV (both present in 48 sequences) upstream of the polyprotein start codon.

We added additional information to our nucleotide alignment: (1) gene annotations (shown as annotation line `#=GC Annotation` in the `stk` file); (2) RNA secondary structures (including pseudoknots) documented in the literature (lines `#=GC SS_cons`, `#=GC SS_cons PK` and the corresponding names specified as Roman numerals in `#=GC SS cons names`); (3) *in silico* predicted novel RNA secondary structures (including alternative conformations), providing a comprehensive view of potential conformations within the *Pestivirus* genome (lines `#=GC SS_cons`, `#=GC SS_cons_alternative` and corresponding names in `#=GC SS_cons names`, `#=GC SS_cons_names_alternative` denoted as ‘SL’ and start position in the genome 1-SD1, accID `NC 076029`); see Tab. 2 and supplementary file F6.

**Table 2:**
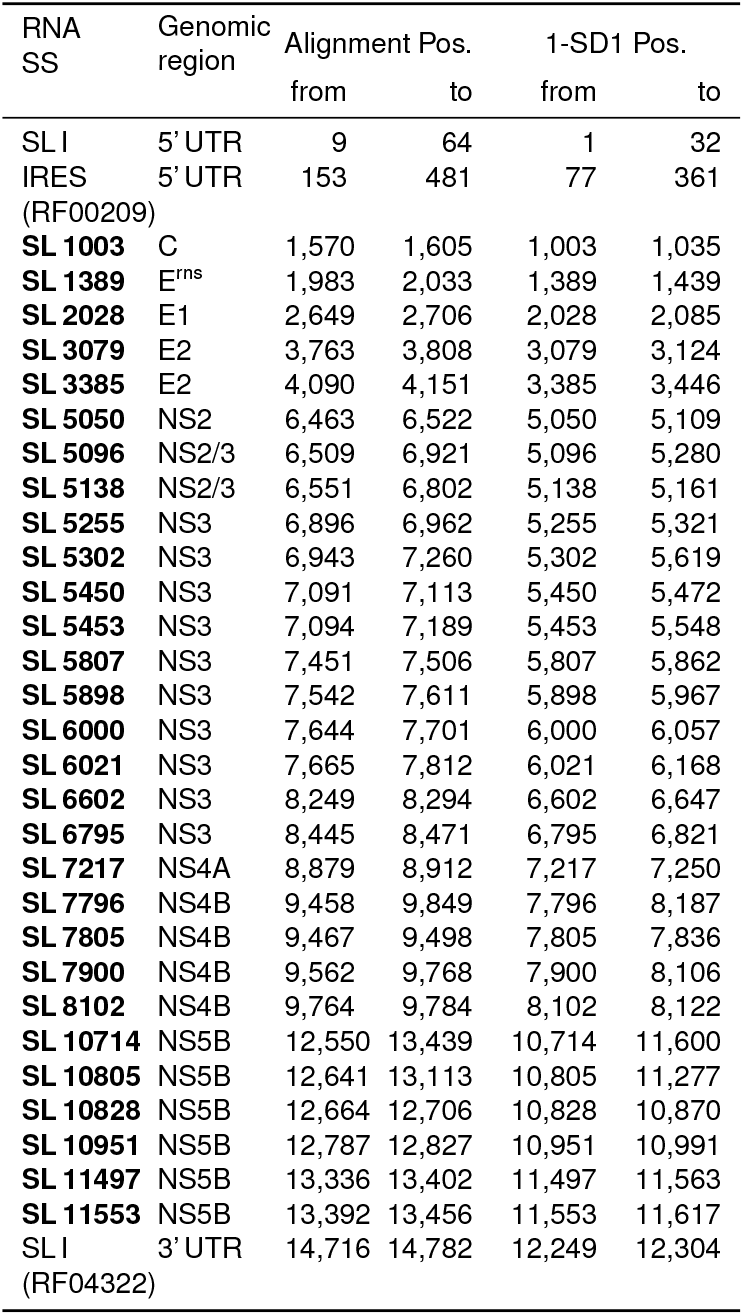
Conserved RNA secondary structures (SS) in *Pestivirus* genome, along with their corresponding `Rfam` model IDs [39, 40]. Start and end positions refer to the alignment and reference strain 1-SD1. We updated the IRES `Rfam` model `RF00209`; confirmed the previously predicted RNA secondary structure of SL I in the 3’ UTR in `Rfam` model `RF04322` (available in `Rfam` v15.1 or later); and predicted further 29 novel conserved RNA families (bold font) which will be incorporated into `Rfam`. We named novel structures according to their start position in the genome 1-SD1 (accID NC_076029).

The protein alignment of the 55 representative genomes encompasses a total of 4,480 residues, with an average of 608 gaps (33,433 gaps in total). We provide the nucleotide and protein alignments in the supplementary material with several formats, such as `Stockholm (stk)`, `ClustalW (aln)`, and `Fasta (fasta)` (see supplementary files F6–F8), which can be conveniently visualized using tools such as `ClustalX` [49, 50], `Jalview` [41] or `Emacs RALEE` mode [51].

### The 5’ UTR of Pestiviruses

The 5’ UTR of pestiviruses consists from 5’ to 3’ end of domain I (D I), a variable region, domain II (D II), and domain III (D III), which is divided into SL IIIa, IIIb, IIIc, IIId_1_, IIId_2_, IIIe, IIIf (also known as stem 1b) and IIIg (also known as stem 1a), see Fig. 3. A pseudoknot (PK, IV) is formed by pairing the region between IIIe and IIIf with the region directly upstream of the start codon, see Fig. 3. D II, D III, and the pseudoknot form together the internal ribosomal entry site (IRES). The alignment of the 5’ UTR is stored in supplementary file F9.

### D I and D II

The very 5’-terminal stem-loop structure of BVDV 5’ UTR has been represented as a bifunctional RNA signal involved in translation and RNA replication [16]. We could identify this domain to be present in all *Pestivirus* species with some expansions of the hairpin loop region, it can be up to 42 nt in length (`NC 077024`, species P).

The variable region between D I and D II harbors for most species a stable hairpin, which is however not at a fixed location but ‘moves’ from species to species up/downstream, as shown previously for `D42109` and `D42110` (both species C) [52]. The D I hairpin also varies in length, shape, internal, and hairpin loops, see Fig. S2.

Three sequences of species C, one of D, and one each of P and Q are predicted to form two hairpins. Some sequences of species A, C, K, and the Zikole strain cannot form a hairpin between D I and D II. Interestingly, despite all the variant RNA secondary structures, the primary sequence is relatively conserved, see Fig. S3. Domain II is a dominant hairpin of high stability in all pestiviruses. Fletcher *et al*. showed that a deletion of nucleotides in D II lead to a reduction of the IRES activity to 19 % [53]. However, the individual length and shape differ, see Fig. S4. The hairpin loop always contains the sequence `CCA` and the 3’ part of the stem consists of the sequence `GUA` in an internal loop in all *Pestivirus* species, suggesting a possible protein interaction, Fig. S5. Domain II was suggested by Gosavi *et al*. to be Y-shaped and supported by SHAPE data in BVDV-1. We could show a potential Y-shaped D II alternative structure for most *Pestivirus* species, however, species P, K, and S can not form a Y-shape. Since these three species are phylogenetically not closely related, we reject a general Y-shape structure of D II in pestiviruses.

### IRES

The IRES of pestiviruses has been analyzed for many years and been well described for decades [52, 54, 55] in BVDV (species A, B) and hog cholera virus (HoCV, nowadays known as classical swine fever virus – CSFV, species C). However, although the `Rfam` DB contains species A–D and G of the IRES, a comparison of species D–S has not been performed yet.

All species share all elements of the IRES, although the sequences differ. However, the RNA secondary structures vary only slightly in the shape and length of the individual hairpins of the IRES. In this study, the IRES element could be computationally retrieved without manual curation with the described methods. Stem IIIg (7 nt) and IIIf (7 nt with one internal loop of 1 nt) are well conserved and completely match in species B, E, and O. Species A, C, D, F, I, L, M, N show a single nucleotide not to basepair, which presumably does not affect the stability of the stem. However, species P, K, Q, R, and J have a disruptive stem IIIg and stem IIIf, which results in a slightly less stable RNA-RNA interaction (ΔMFE = 1.5 kcal/mol). Stem-loop IIIa has in each species exactly the length of 6 nt with a hairpin loop length of 4 not, indicating the evolutionary constraints of that hairpin. The strongly conserved consensus motif for the hairpin loop and closing base pair is `GAGUAC`. Hairpin IIIb varies in length (16-25 nt) and stability (MFE ranging from −13.50 to - 27.60 kcal/mol). The hairpin loops of species K and S are longer (25 nt compared to usually 19 nt). Stem IIIc is highly compensatory mutated however, it maintains in each species the same length of 3 nt, with a variant hairpin loop of 3 (species Q, R, and J) or 4 nucleotides (other species), which is a conserved motif in species A, B, and C, divergent for other species. Stem IIId_1_ is in each species exactly 10 nt long with a hairpin loop of 4 or 5 nucleotides containing always the sequence `GGGG`. Stem IIId_2_ differs from species to species in sequence, length (3-11 nt) and hairpin length. Stem IIIe consists of 4 base pairs (or 3 bp in some variants of species C) and 4 nt in the loop region.

### Pseudoknot

The IRES pseudoknot is in each species very stable (MFE ranging from −6.20 to −10.60 kcal/mol). Some publications suggest another stem-loop of length two surrounding the pseudoknot interacting part between stem IIIe and IIIf [53, 54]. However, this can not be confirmed for most of the species and has recently also been shown by SHAPE data to be not present for BVDV [13]. We propose instead an extended, slightly more stable pseudoknot interaction of 8 nt instead of only 6 nt, as also previously suggested by SHAPE data [13], see Fig. 4 (ΔMFE=1.63 kcal/mol).

**Figure 4:**
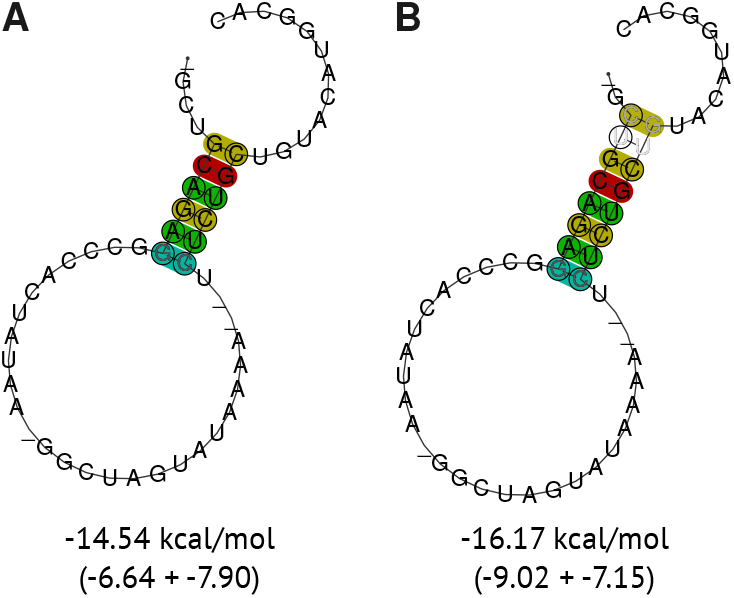
Consensus RNA secondary structure of the pseudoknot in the 5’ UTR of the (A) 6 nt stem-loop and (B) 8 nt stem-loop. The structures were predicted and visualized using `RNAfold` with additional parameter for structural constraints. The constraints forced the lengths of the stem-loops. The averaged MFE in kcal/mol (composed of actual MFE and covariance pseudo-energy) are displayed below the structure.

Another hairpin has been discussed several times directly upstream of the start codon [53]. We could not find this hairpin conserved for all pestiviruses. We hypothesize this hairpin to be present for all pestivirus species A, B, and C, as well as in some other pestivirus isolates (G, H, D, I, N, O, and E). However, its functionality remains debated, because this hairpin competes with the formation of the pseudoknot.

### Other observations in the 5’ UTR

The palin-dromic regions V1, V2, and V3 reported in previous studies [56–58] correspond to stem IIIb, IIId_1_, and IIId_2_, respectively, and have been used for genotyping.

In the work of Giangaspero *et al*. [57] claimed for species D a phylogenetic reconstruction being possible only based on the 5’ UTR. We generated a phylogenetic representation using `SplitsTree` for 51/55 pestivirus genomes (see Fig. S1, including all genomes with a present 5’ UTR) and could confirm a high similarity of the split graph based on only the 5’ UTR of pestiviruses compared to the tree based on entire genomes.

### The 3’ UTR of Pestiviruses

The 3’ UTR of pestiviruses has been shown mainly for BVDV and CSFV to consist of stem-loop (SL) IV, III, II, and I [55, 59]. A comprehensive analysis of the 3’ UTR across several hundred *Pestivirus* sequences has not yet been performed. Within our analysis, only SL I is conserved across all *Pestivirus* species, see Fig. 5 and supplementary file F10. Although SL I shows sequence divergence outside of pestivirus species A, B, and C, it still forms a conserved secondary structure, as shown in Fig. S6. According to a study by Pankraz [18], the presence of SL I and the 3’-terminal region between SL I and II are crucial for pestivirus replication. In contrast, deletions within SL II and SL III, or even the absence of either of these loops, did not significantly affect viral replication. However, a mutant RNA lacking both SL II and SL III was found to be non-infectious. This suggests that while SL II and SL III are less critical individually, the presence of at least one of these elements was shown to be essential for the replication of BVDV-1 strain CP7.

**Figure 5:**
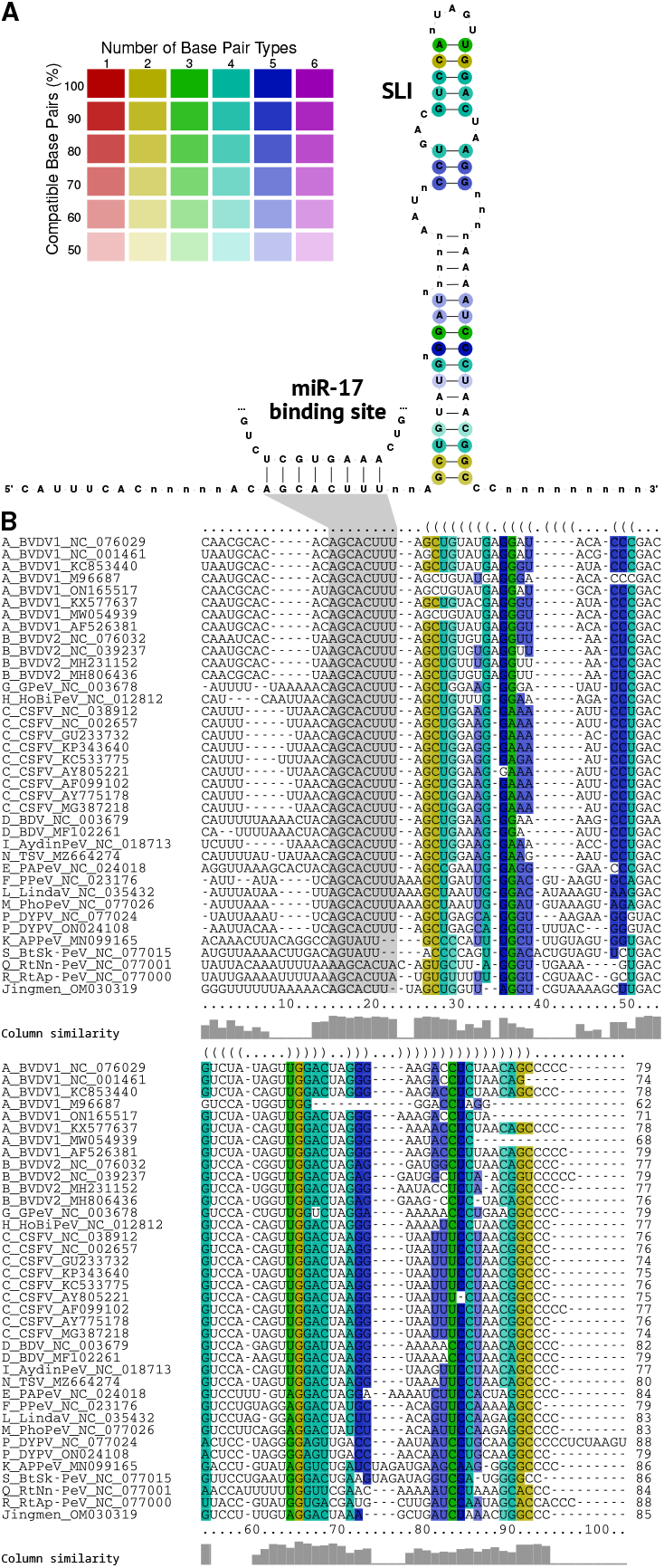
(A) RNA secondary structure of the 3’ UTR SL I, colored by the number of base pair types and the percentage of sequences that form the base pairs, illustrating the extent of covariations in double-stranded regions. (B) Multiple sequence alignment of the 3’ UTR SL I. Among the selected sequences, 38 isolates had a 3’ UTR sequence in the region of SL I. The binding site for miR-17 is located upstream of the stem-loop with the seed sequence AGCACUUU present in 33 sequences (highlighted in gray).

### Species A and C

Further structures in the 3’ UTR (SL II, SL III, SL IV) were reported for the species A–C [17, 18, 55]. However, the 3’ UTR region is very heterogeneous [45], as the 3’ end of S IV in species A appears homologous to the 5’ end of S III in species C. Therefore, SL II appears larger in species A than in other species. Subsequently, SL III can be formed only for longer sequences (`NC 076029`, `NC 001461`, `ON165517`) of species A, as previously reported [17, 55]. Species-specific alignments for the 3’ UTR are stored in supplementary files F11 and F12.

### SL IV

Generally, SL IV is present in all sequences of species A, however, it is difficult to reconstruct the evolution of SL IV. Our prediction of SL IV in species C slightly differs from the literature [55], Fig. 6.

**Figure 6:**
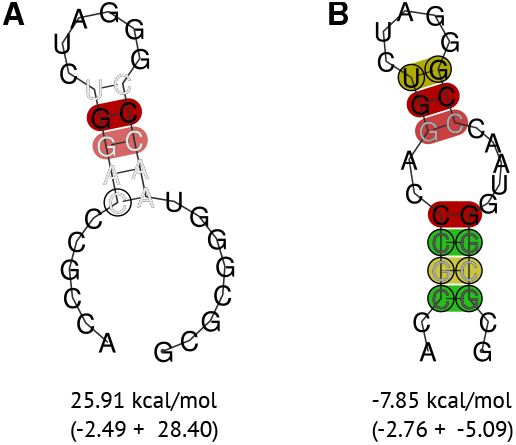
SL IV of species C. The previously reported (A) secondary structure [55] can only be formed in 4 out of 10 representative C sequences (`GU233732`, `KP343640`, `KC533775`, `MG387218`), whereas our predicted secondary structure of SL IV (B) can be formed by all sequences with MFE ranging from −8.10 to −13.50 kcal/mol. The averaged MFE in kcal/mol (composed of actual MFE and covariance pseudo-energy) are displayed below the structure.

### AU-rich region

SL III has been reported to be AU-rich in strain C [45, 60]. In the present work, we additionally find species D, I, N, O, E, and L to contain such an AU-rich region. Interestingly, species E and L are phylogenetically distant from D, I, N, and O (see Fig. 1=, and we hypothesize the AU-rich region to be invented twice during *Pestivirus* evolution. Additionally, species F is phylogenetically related to species E and L, but appears without an AU-rich region.

### MicroRNA binding sites

MiR-17 is known to bind to the 3’ UTR of pestiviruses in species A, B, and C [20, 61]. The binding of miR-17 to the 3’ UTR is essential for replication, enhances viral translation, and increases RNA stability. In the present work, the seed sequence (AGCACUUU) of the miR-17 binding site directly upstream of SL I is perfectly conserved in 33 sequences from 13 species covering that region. Species K, S, Q, R, and the Jingmen strain show mutations and deletions in the seed sequence, see Fig. 5, thus we claim no interaction with miR-17 to take place. No sequences of species O and J were available for that region.

The binding of microRNA let-7 is known to increase viral translation to its full efficiency [20]. In species A and C, the binding site is located in the hairpin loop of SL II. The seed sequence CUACCUCA is located further upstream of the miR-17 binding site. Here, we identified 38 representative sequences from 11 species (A-D, F-I, L, N, O) that show a strong conserved binding site of let-7.

### Other conserved regions in specific species

All other sequences only show similarities within species or some conserved region at the primary sequence level between some species. The sequences of the species K are very divergent but are generally alignable at a sequence level, see supplementary file F13. Species K and S show fragment-wise sequence similarity. When comparing species K, S, and Zikole, we observe primary sequence similarities in the location of SL III (alignment pos. 14,151 to 14,245). Still, we could not predict a conserved secondary structure for all three species. The closely related species Q, J, and R contain the longest sequences within the 3’ UTR, with an insertion upstream of SL I, which shows sequence similarities between the three sequences. The three representative sequences of species P are generally well conserved.

### New RNA secondary structures candidates in the protein-coding region

The importance of RNA secondary structures in the protein coding region and the underlying selection pressure have been extensively studied, showing that structural conformations in the CDS can be beneficial for viral translation and replication [62, 63]. Prediction of RNA secondary structures based on alignments was performed in a previous study for HCV and resulted in over 40 structural elements (including 23 novel ones) in the coding region [64]. Although the computational method and overall genome organization are similar here, the results should not be compared because HCV is a species and pestivirus is an entire genus with greater sequence diversity.

Our analysis identified several novel RNA secondary structures in the protein-coding region using our *in silico* prediction method, see Tab. 2, Fig. 7, and S7. We selected new structure candidates based on the conservation at both the sequence and structural levels, particularly the presence of compensatory mutations. We identified 29 novel structures that are likely functional elements, as they exhibit conserved RNA secondary structures with compensatory mutations despite being located in coding regions (Fig. 7 and S7). Furthermore, we identified three structures that are conserved in a subset of species (see Tab. S3).

**Figure 7:**
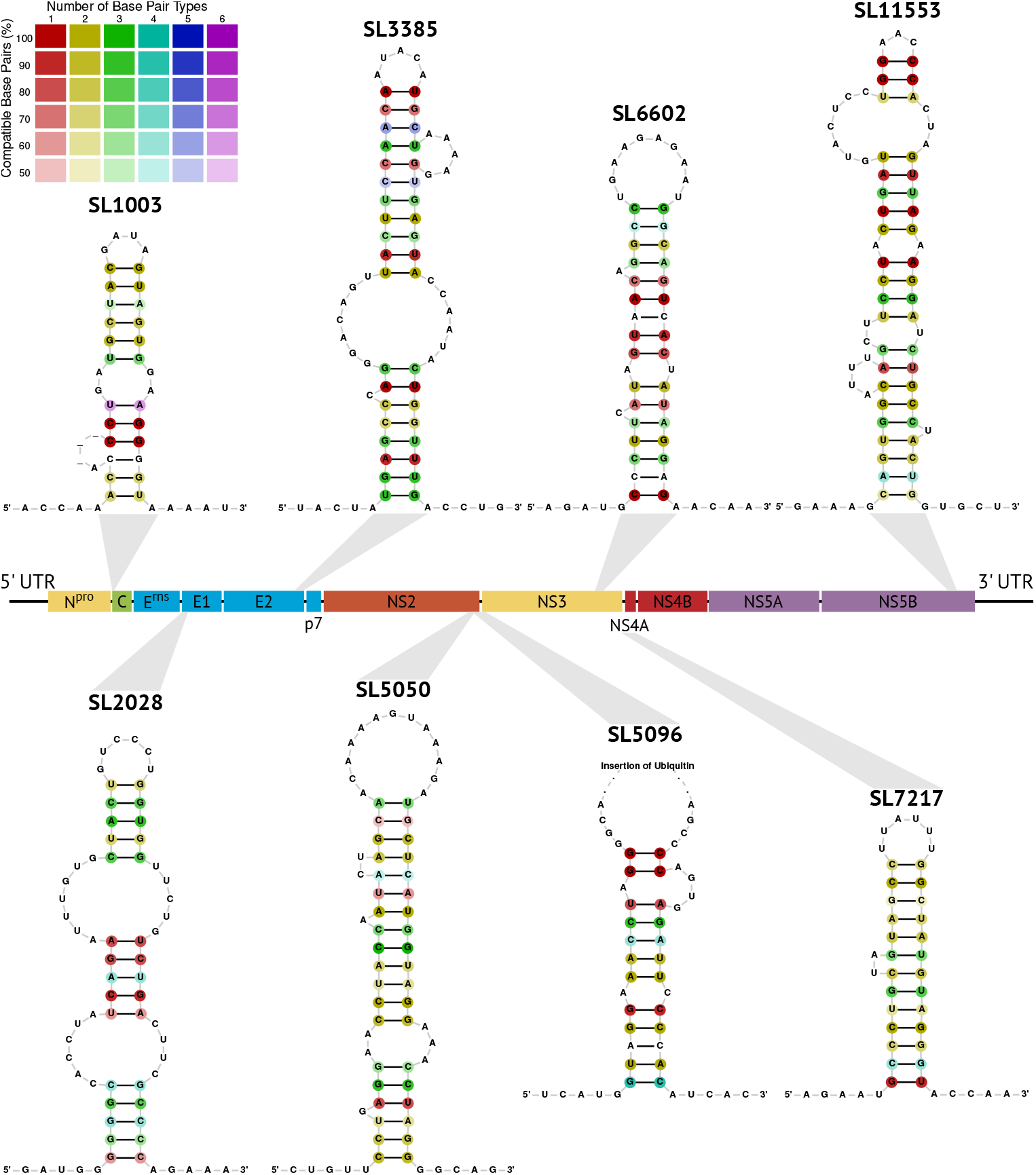
Selection of eight novel conserved RNA secondary structure candidates located in the protein-coding region of the *Pestivirus* alignment. Novel structures are named according to their start position in the genome 1-SD1 (accID `NC_076029`).

These well-conserved structures represent previously unrecognized RNA elements within representative *Pestivirus* genomes, highlighting the complexity and diversity of RNA secondary structures in this viral genus. The discovery of these novel structures provides new insights into their potential roles in *Pestivirus* replication, translation, and pathogenesis. Further investigations are necessary to experimentally validate their function and explore their biological significance in the context of *Pestivirus* infections.

### Improvements to Rfam Virus Families

In total, we identified 32 conserved structural regions across the representative genomes of pestiviruses, of which 29 are presented here for the first time. Only the 5’ UTR of the pestiviral genome is currently present as a known RNA secondary structure in `Rfam`. We improved the IRES model (`RF00209`) from the `Rfam` database [39, 40] (see Tab. 2) to ensure comprehensive coverage of the entire phyloge-netic clade of the pestivirus sequences. The previous IRES model contained 25 sequences only covering the species A, B, C, D, and G. We extended the model to 48 sequences covering all 19 species, spanning 508 nucleotide positions in the alignment (previously 286). The new model now includes SL I (from 32 *Pestivirus* genomes), which was absent due to sequencing problems of the very 5’ genome end in previous times. Importantly, the new model now includes the pseudoknot SL IV. In addition, we provide alignments for the new models of RNA secondary structures in the 3’ UTR for the `Rfam`. `RF04322` includes SL I, which is conserved throughout the entire genus of pestiviruses (see Tab. 2 and Fig. 5), while the models `RF04323`, `RF04324` and `RF04325` represent species A, C, and K, respectively (see Tab. S3).

### Genome circularization

Genome circularization is a well-known phenomenon, especially for the members of the genus *Orthoflavivirus*, and is considered important for viral replication and drug design [65–67]. In this study, we predicted a potential long-range interaction (LRI) between the 5’ and 3’ UTRs in 29 of our representative *Pestivirus* sequences (15 species), where both UTRs are present (31). Typically, genome circularization involves interactions between the very 5’ end of the 5’ UTR and the 3’ end of the 3’ UTR. However, our prediction suggests that the LRI forms upstream of the polyprotein start codon and the very 3’ end of the 3’ UTR, see Fig. 8. This implies that SL IIIc to IIIe in the 5’ UTR and SL I in the 3’ UTR must unfold to allow the LRI to form. While our findings provide new insights into possible genome circularization mechanisms in *Pestivirus*, they remain based solely on *in silico* predictions and require experimental validation. Experiments demonstrating that the pestiviral IRES element can be functionally replaced by IRES sequences from HCV or even encephalomyocarditis virus (EMCV) argue at least against an essential role of these interactions [68].

**Figure 8:**
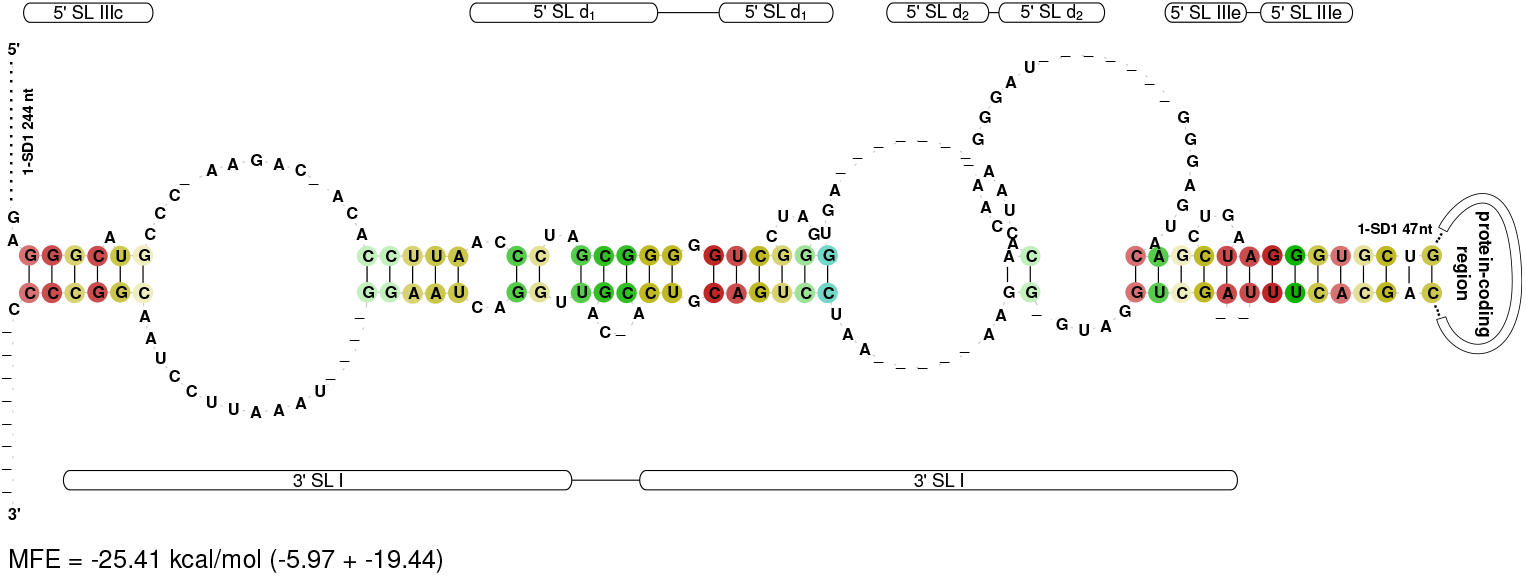
*In silico* predicted RNA-RNA interaction between the 5’ and 3’ UTRs of 29 representative *Pestivirus* genomes (15 species). The possible genome circularization is located upstream of the start codon and at the very 3’ end of the genome with an MFE of −25.41 kcal/mol.

### Larger host insertions

Our alignment captures representative pestiviral genomes with insertions from host mRNA, which are reported features of cytopathogenicity. Insertions are typically found in cytopathogenic pestiviruses [11, 12, 69, 70]. The insertion of DNAJC14 (nt aln pos. 6,017–6,398, aa aln pos. 1,836–1,964) is present in the following three genomes of our representative set, see Fig. 2B orange box and Fig. 9A: strain NADL (species A, accID `NC_001461`), isolate 296c (species B, accID `MH806436`) and giraffe-1 H138 (species G, accID `NC_003678`). DNAJC14 is incorporated into the genome of pestiviruses within the NS2 gene (see introduction). The Jingmen isolate SYS_SheQu (species unclassified, accID `OM030319`) also contains an insertion, which is, however, not homologous to DNAJC14 and also seems not to match to any known sequence in the NCBI. Another insertion is known between the genes for NS2 and NS3. This is the ubiquitin insertion (nt aln pos. 6,563–6,793, aa aln pos. 2,021–2,095), which is only present in one of our representatives; the Osloss strain (species A, accID `M96687`), see Fig. 2B cyan box and Fig. 9B. Other insertions, such as NEDD3, SUMO, and LC3 [12], are not included in the MSA, as these sequences are only part of partial genomes and therefore do not occur in our original data set. We expect that extensive sequencing in the next few years will lead to genome sequences with such insertions that can be included in our alignment. Our results show that our method can preferentially select representative genomes that are divergent in terms of insertions.

**Figure 9:**
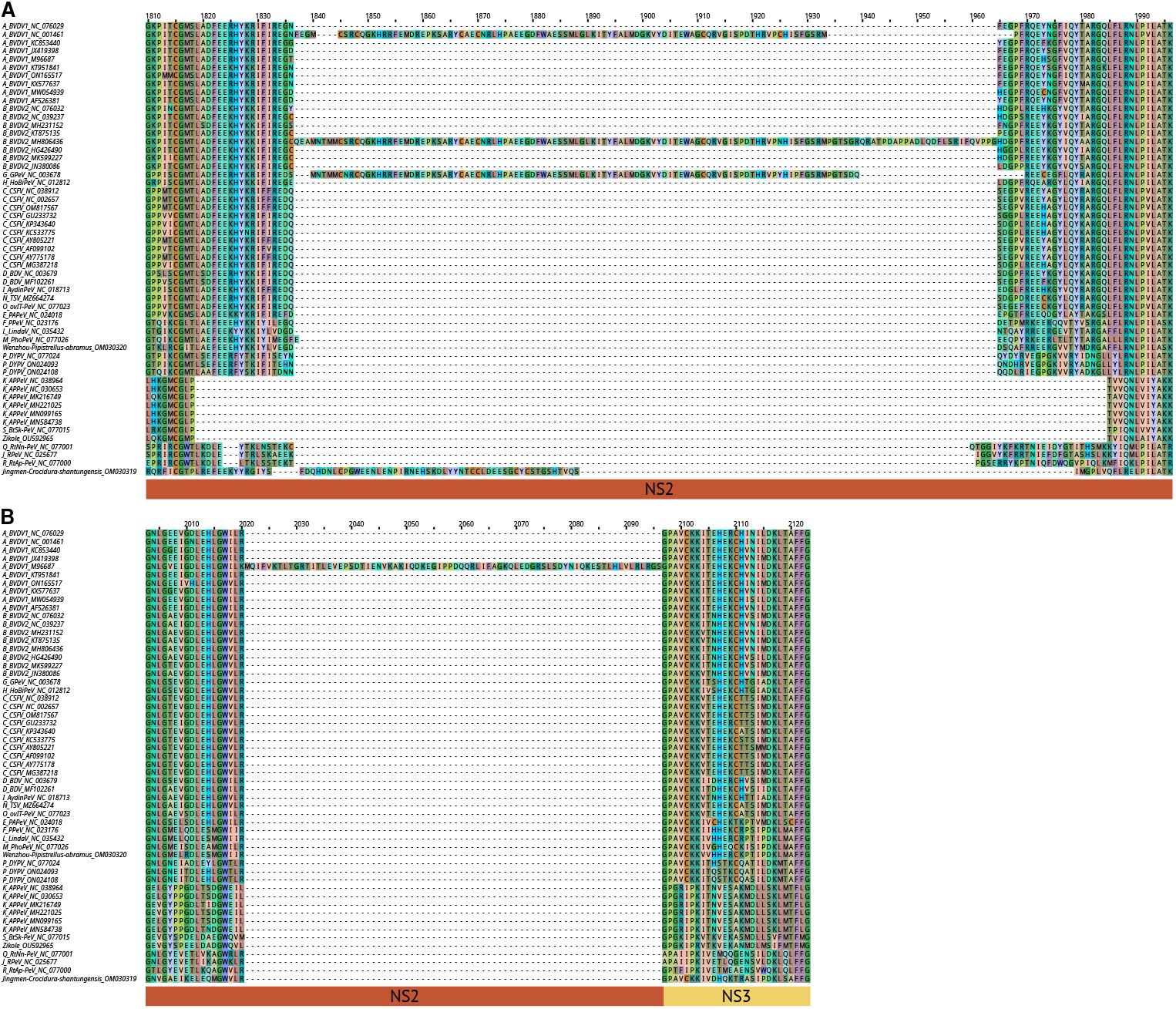
Multiple sequence alignments showing the insertion of (A) DNAJC14, see Fig 2 orange box and (B) ubiquitin, see Fig. 2 cyan box. (A) The insertion is of DNAJC14 present in the NADL strain (species A, accID `NC_001461`), isolate 296c (species B, accID `MH806436`), and giraffe-1 H138 (species G, accID `NC_003678`). An undescribed insertion occurs in the Jingmen isolate SYS SheQu (species unclassified, accID `OM030319`). (B) The insertion of ubiquitin is only present in the Osloss strain (species A, accID `M96687`).

### Potential Primer Sites for Pestivirus Detection

Using our tool `AnchoRNA`, we identified a set of highly conserved regions at the nucleotide level across pestivirus sequences. These regions represent promising candidates for primer design and may serve as molecular markers for pestivirus detection via PCR (see Fig. S8). The corresponding list of conserved anchors for each sequence is summarized in Tab. 3. The table shows how many of the 55 representative pestivirus sequences contain each conserved region, indicating their broad applicability across pestivirus species. We emphasize that the proposed primers have not been experimentally validated. Rather, we present these conserved regions as potential starting points for the design and evaluation of molecular diagnostics. Experimental testing is required to assess their sensitivity, specificity, and overall performance.

**Table 3:** Detection of representative pestivirus sequences (out of 55) using primers derived from selected anchors for pestivirus identification. This table holds all potential primer sites for at least pestiviruses A, B, and C, see Fig. S8.

|  |  |  |  |  |  |  |  |  |  |  |  |
| --- | --- | --- | --- | --- | --- | --- | --- | --- | --- | --- | --- |
| Anchor No. | 12 | 14 | 34 | 38 | 39 | 59 | 71 | 72 | 74 | 77 | 81 |
| #Seq | 53 | 46 | 43 | 44 | 38 | 39 | 46 | 52 | 51 | 45 | 38 |

### Jingmen *Crocidura shantungensis* genome

The Jingmen sequence (isolate SYS SheQu, accID `OM030319`) has been reported to be a *Pestivirus* 1 isolate in the NCBI database (accessed March 17, 2025). Although some of the protein-coding regions are alignable with high conservation to *Pestivirus* species (protein C, NS3, NS4A, NS4B, NS5B) other regions are on nucleotide and amino acid levels harder to align computationally or manually (protein N^pro^, E^rns^, E1, E2, NS5A). Protein NS2 seems to have very little similarity to other *Pestivirus* species. The non-essential region (directly downstream of the 5’ UTR) is very short (similar to only species M). The supposed 5’ UTR contains neither any single RNA secondary structure element of the IRES, or D I, nor any conserved motif, which leads us to the conclusion, that this region is (a) either a reminent of the non-essential region (however, it is not translatable into a protein) and the 5’ UTR of the Jingmen sequence is absent, (b) a chimeric sequence is present, or (c) it might be a sequencing artifact. For this reason, we decided to truncate the Jingmen sequence upstream of the start codon.

## Conclusion

In this study, we present the first comprehensive genome-wide multiple sequence alignment (MSA) of pestivirus genomes, incorporating computational predictions of RNA secondary structures. Our approach leveraged a representative selection of 55 genomes, chosen through `HDBSCAN` clustering of k-mer distributions and dimension reduction, effectively capturing the phylogenetic and functional diversity of the genus, e.g., host mRNA insertions. The alignment integrates existing genome annotations, conserved structural elements, and pseudo-knots, facilitating seamless comparison with other research studies. By incorporating sub-optimal structure predictions via tools such as `LocARNA` and `RNAalifold`, we reveal potential alternative RNA conformations that may play regulatory roles. These findings expand our understanding and provide new insights into the viral ‘life cycle’ of pestiviruses.

While our study offers valuable insights into conserved RNA secondary structures in pestivirus genomes, it has limitations. It does not automatically account for long-range RNA–RNA interactions across the whole genome, three-dimensional RNA conformations, or RNA–protein interactions, all of which may play critical regulatory roles and warrant further investigation. The computational challenges associated with modeling such long-range interactions remain unresolved and have been tackled only partially in the past, as for HCV [71]. In addition, future studies should aim to validate the predicted structures and their biological relevance *in vivo*, identify other potentially involved miRNAs, and assess the functional implications of these conserved elements.

All conserved RNA secondary structure models identified here will be included in `Rfam` version 15.1 or later, ensuring broad accessibility and utility. This alignment provides a standardized and extensible framework for future pestivirus studies, serving as a foundational resource for comparative genomics, evolutionary analyses, and functional investigations.

## Authors contributions

Sandra Triebel: Investigation; Data curation; Methodology; Formal analysis; Visualization; Writing – original draft; Writing – review & editing. Tom Eulenfeld: Software; Writing – review & editing. Nancy Ontiveros-Palacios: Data curation; Writing – review & editing. Blake Sweeney: Data curation; Writing – review & editing. Norbert Tautz: Validation; Writing – original draft; Writing – review & editing. Manja Marz: Supervision; Conceptualization; Investigation; Data curation; Methodology; Formal analysis; Writing – original draft; Writing – review & editing.

## Availability of data and materials

Data is available in the supplementary information and via Zenodo https://doi.org/10.5281/zenodo.15490752.

The RNA secondary structure models are provided in the `Rfam` database [39, 40].

## Code availability

`ViralClust` is available via GitHub https://github.com/rnajena/viralclust [23]. `AnchoRNA` is available via GitHub https://github.com/rnajena/anchorna [36]. `sugar` is available via GitHub https://github.com/rnajena/sugar [43].

## Ethics approval

Not applicable

## Consent to participate

Not applicable

## Consent for publication

Not applicable

