## Supplementary Information for "First full-genome alignment representative for the genus *Pestivirus*"

### Files

- F1 - original data set in `Fasta (fasta)`
- F2 - pre-filtered data set in `Fasta (fasta)`
- F3 - phylogenetic tree in `Newick (nwk)` and `Nexus (nex)`
- F4 - representative genomes in `Fasta (fasta)`
- F5 - `AnchoRNA` output in `General feature (gff)` (nucleotide and protein sequences)
- F6 - nucleotide alignment in `ClustalW (aln)`, `Fasta (fasta)`, and `Stockholm (stk)` (latter with additional annotations such as RNA secondary structures and genes)
- F7 - nucleotide alignment of the CDS in `ClustalW (aln)`, `Fasta (fasta)`, and `Stockholm (stk)` (latter with additional annotations such as RNA secondary structures and genes)
- F8 - protein alignment in `ClustalW (aln)`, `Fasta (fasta)`, and `Stockholm (stk)`
- F9 - nucleotide alignment of the 5' UTR (51 sequences) in `Stockholm (stk)`, `ClustalW (aln)`, and `Fasta (fasta)`
- F10 - nucleotide alignment of the 3' UTR (52 sequences) in `Stockholm (stk)`, `ClustalW (aln)`, and `Fasta (fasta)`
- F11 - nucleotide alignment of the 3' UTR of species A (10 sequences) in `Stockholm (stk)`
- F12 - nucleotide alignment of the 3' UTR of species C (10 sequences) in `Stockholm (stk)`
- F13 - nucleotide alignment of the 3' UTR of species K, S, and the Zikole strain (7 sequences) in `Stockholm (stk)`

### Clustering

**Table S1:** Clustering results using `ViralClust`. # cluster – number of clusters (with a size greater than one); min. size – size of smallest cluster; max. size – size of largest cluster; avg. size – average cluster size; median size – median cluster size; d. centroid – average phylogenetic distance between two representative genomes; unclustered – number of unclustered sequences (= cluster with only one element or determined as unclustered by `HDBSCAN`); cps – average cluster per species; cpg – average cluster per genus. The last two pieces of information give a measurement of how much a single species/genus is divided into different groups.

| algorithm | # cluster | min. size | max. size | avg. size | median size | unclustered | d. centroid | cps | cpg |
|---|---|---|---|---|---|---|---|---|---|
| `cd-hit-est` | 68 | 2 | 98 | 10.10 | 3 | 69 | 1.19 | 2.46 | 41.0 |
| `HDBSCAN` | 53 | 6 | 40 | 12.89 | 10 | 73 | 0.98 | 2.39 | 25.5 |
| `MMSeqs2` | 65 | 2 | 106 | 10.55 | 3 | 70 | 1.28 | 2.24 | 37.0 |
| `sumaclust` | 63 | 2 | 101 | 9.86 | 4 | 56 | 1.34 | 2.33 | 43.0 |
| `vclust` | 64 | 2 | 107 | 10.98 | 3 | 53 | 1.27 | 2.03 | 42.0 |

### Representative genomes

**Table S2:** Selection of representative genomes.

| | |
|---|---|
| RefSeqs | NC_024018, NC_023176, NC_018713, NC_076029, NC_001461, NC_012812, NC_003678, NC_035432, NC_003679, NC_077024, NC_025677, NC_038964, NC_038912, NC_076032, NC_039237, NC_077000, NC_077001, NC_077015, NC_077023, NC_077026, NC_030653, NC_002657, MZ664274 |
| Reduce overrepresentation | KT875160, AB894423, HQ174297, MH231142, LC649064, LT158404, JX297520, HQ174294, OR004808, MH221026, KC695814, HM237795, MH885413, MT799517, LT593753, MK121886, MK093249, MN558876, MW528229, JQ612704, KJ463422, AJ133738, AF091605 |
| Outlier | MW256672, MN025505, ON684360, OM030319, JQ799141, OU592965, OM030320 |
| Highly similar sequences (temporarily removed) | NC_076029, KC853440, JX419398, KT951841, ON165517, KX577637, AF526381, NC_076032, NC_039237, MH231152, KT875135, HG426490, MK599227, NC_038912, NC_002657, GU233732, KP343640, KC533775, AY805221, AF099102, AY775178, NC_038964, MH221025, MN099165, MN584738 |

### 5' UTR

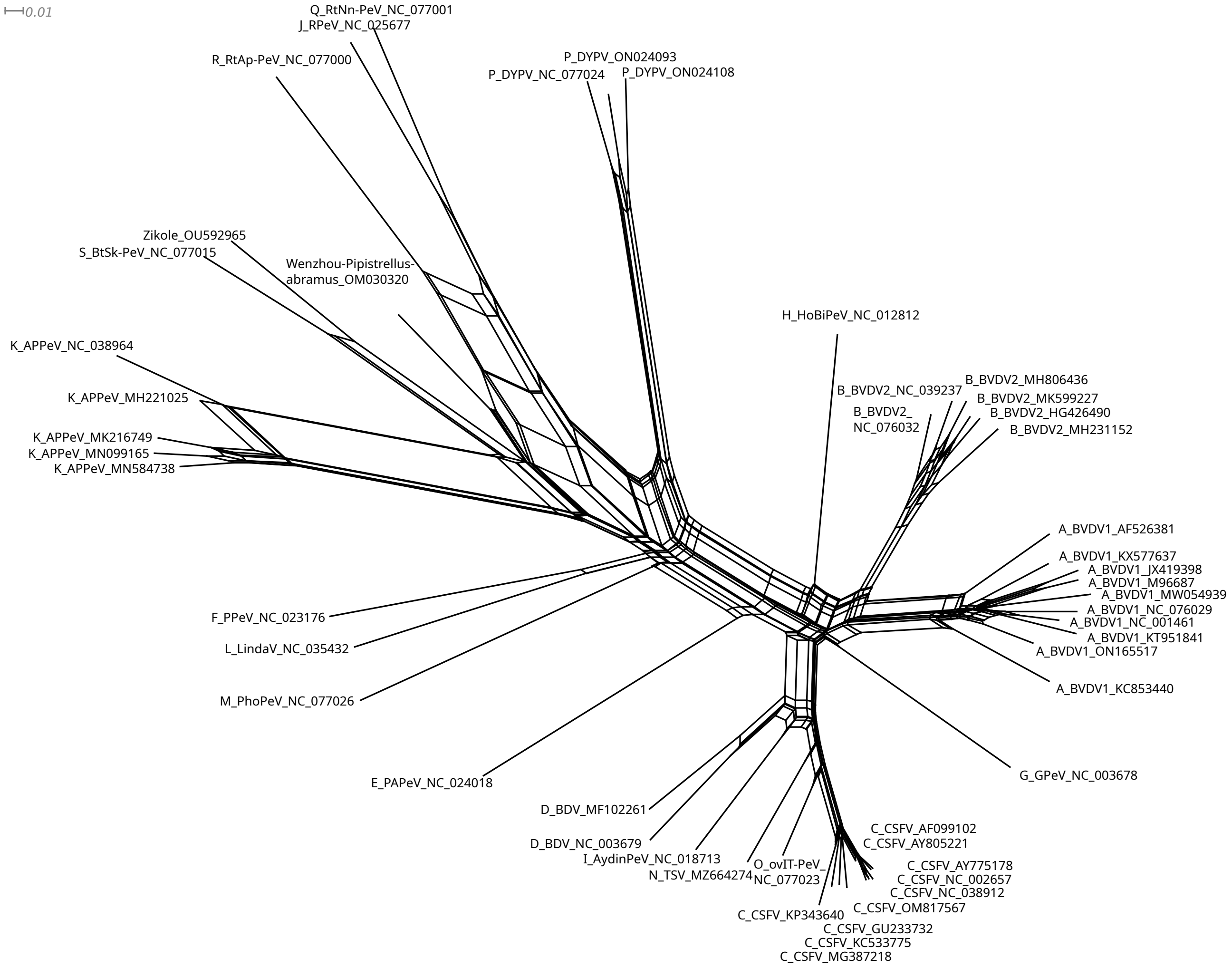


**Figure S1:** Phylogenetic reconstruction based on the 5' UTR alignment. The tree is based on the `MAFFT` alignment of the 5' UTR sequence of the representative genomes (if present) and served as input for `SplitsTree` to construct a splits graph using the neighbor-net algorithm. The overall structure of the tree stays the same compared to the whole-genome phylogeny, see Fig. 1. Isolates of one species cluster together in one subtree, e.g., A, B, C. Species G and H are not clustered right next to each other, but still close. Species F, L, M, and E are still close to the root but located on the other side of the tree in this visualization. However, the placement of Wenzhou is different; instead of being between species F and L, it is now between species S and R.

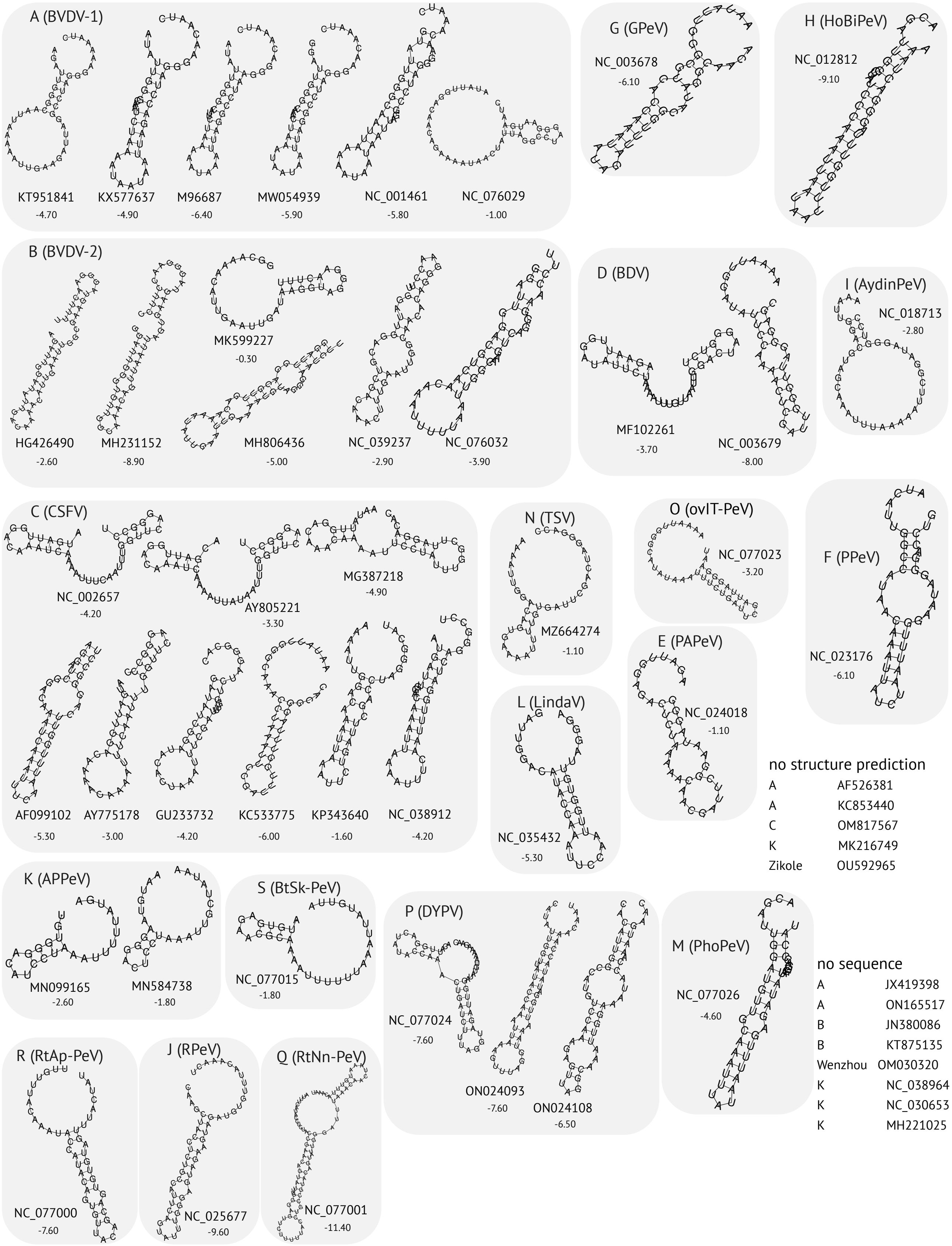


**Figure S2:** Single-sequence RNA secondary structure prediction of the region between SL I and SL II in the 5' UTR. MFE in kcal/mol of each structure is displayed below the Genbank ID.

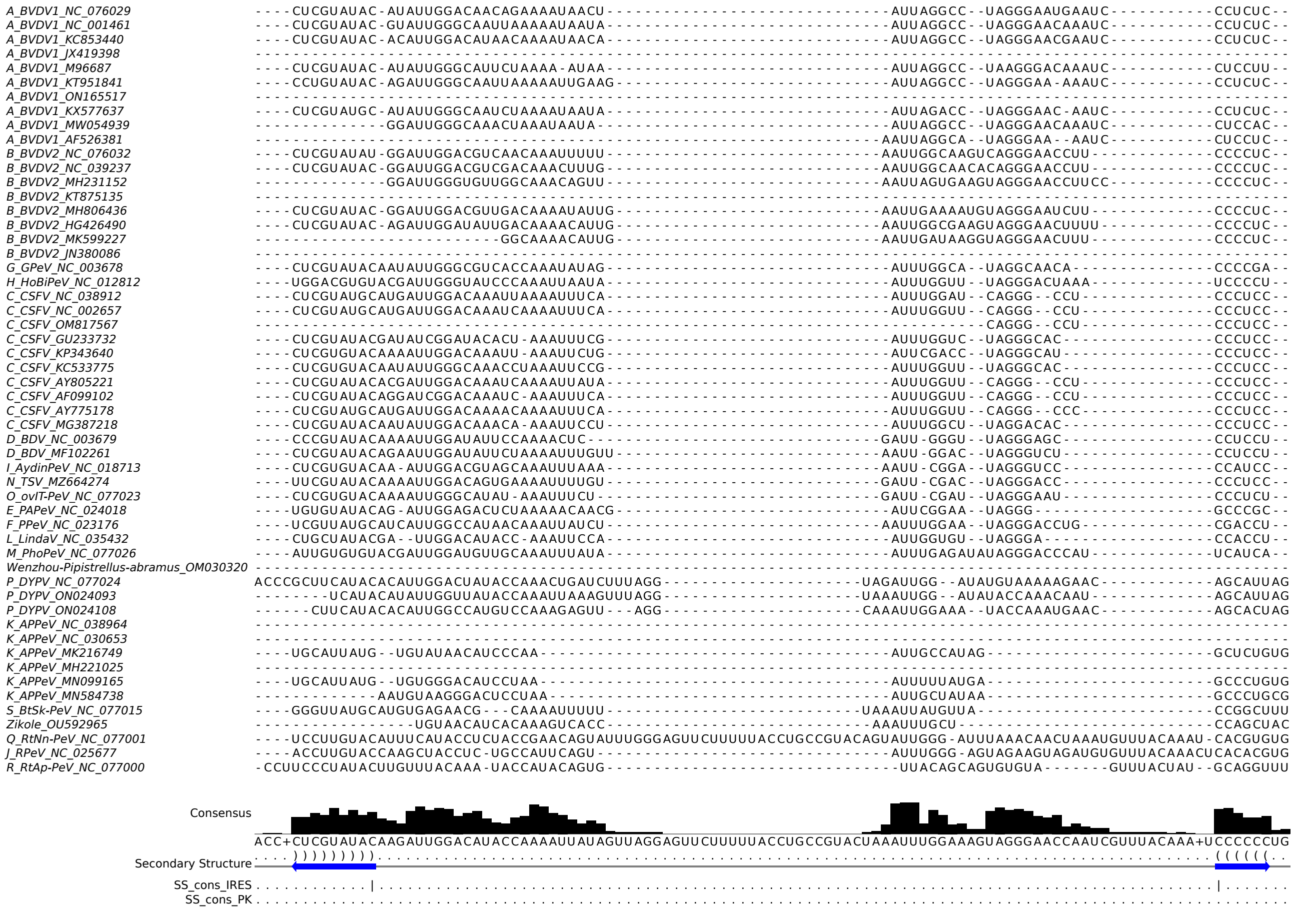


**Figure S3:** Multiple sequence alignment of the region between SL I and II in the 5' UTR (see Fig. 3 and S2).

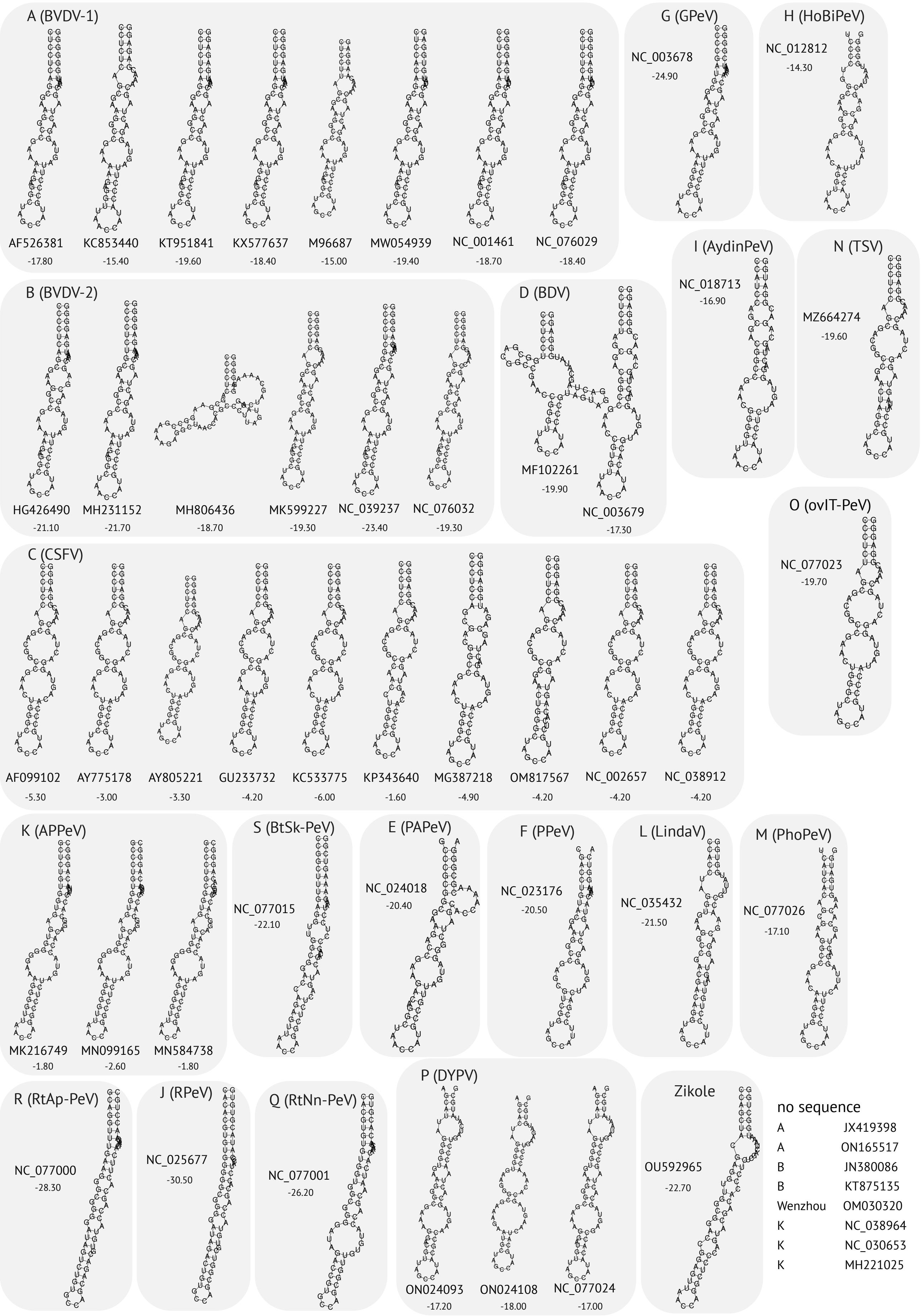


**Figure S4:** Single-sequence RNA secondary structure prediction of SL II in the 5' UTR using `RNAfold` with additional parameter for structural constraints forcing the four nucleotides in the hairpin loop to be unpaired. MFE in kcal/mol of each structure is displayed below the Genbank ID.

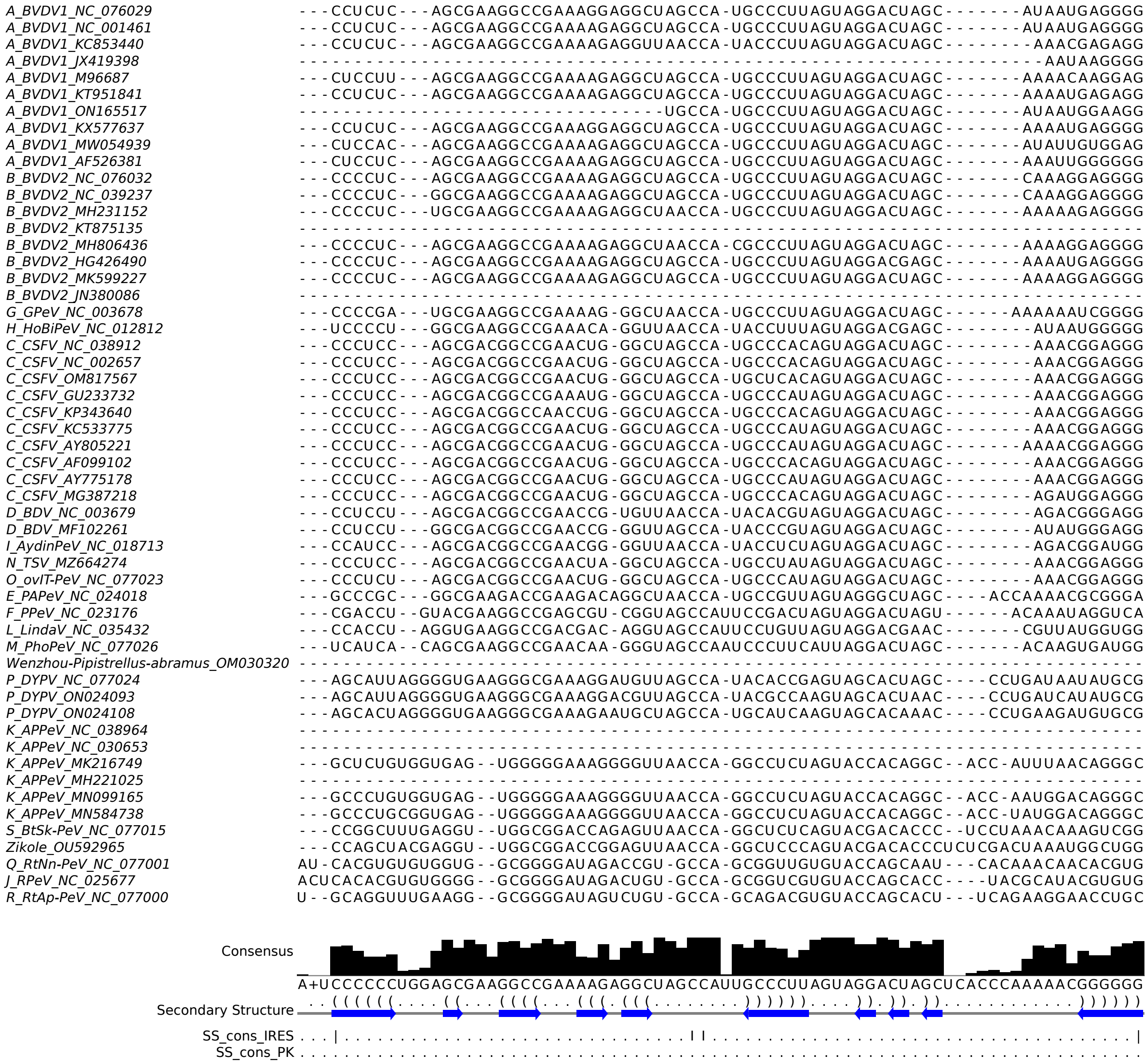


**Figure S5:** Multiple sequence alignment of SL II in the 5' UTR (see Fig. 3 and S4).

### 3' UTR

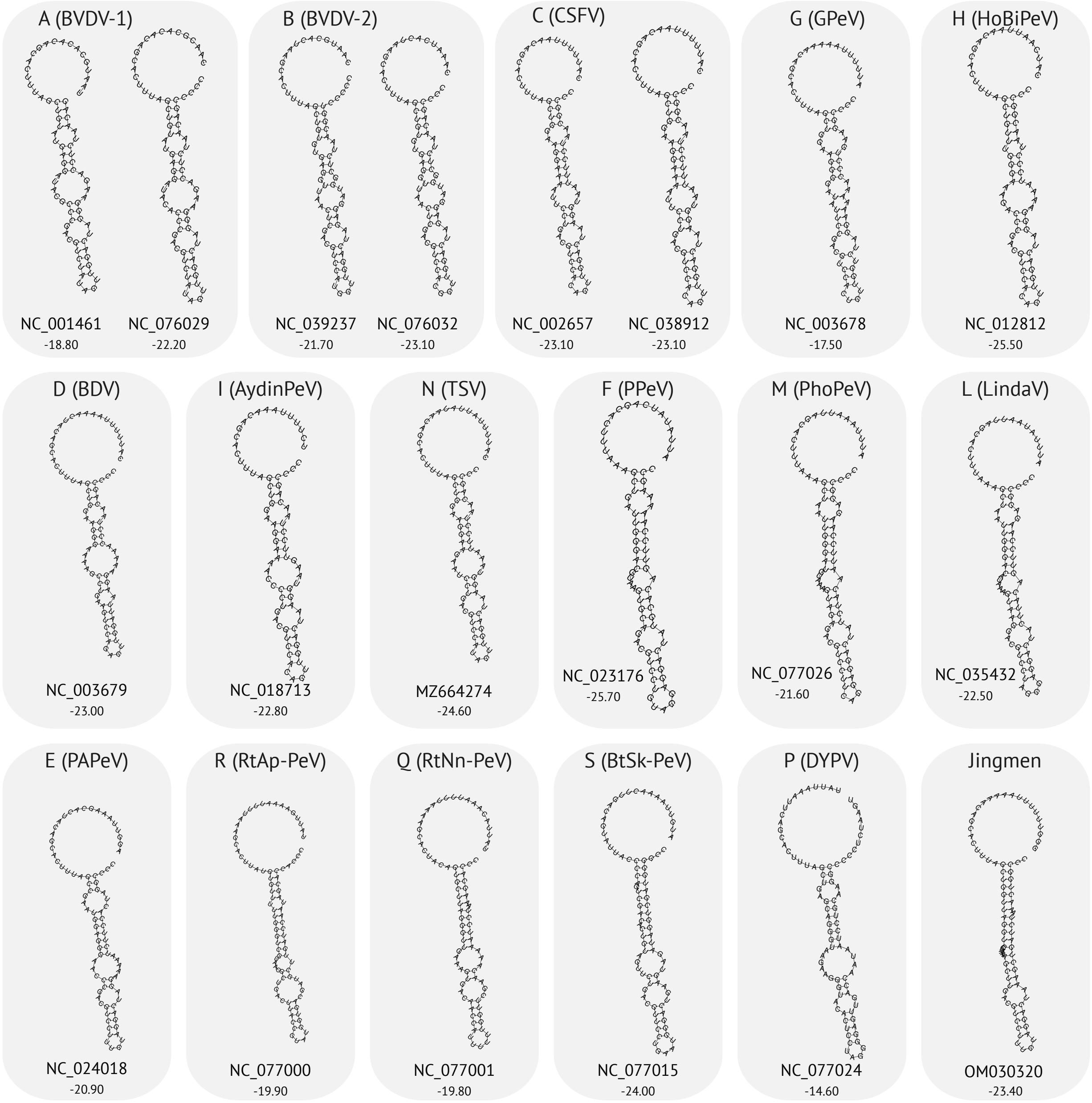


**Figure S6:** Single-sequence RNA secondary structure prediction of SL I in the 3' UTR using `RNAfold` with additional parameter for structural constraints forcing the miR-17 biniding site to be unpaired. MFE in kcal/mol of each structure is displayed below the Genbank ID.

### New structure candidates

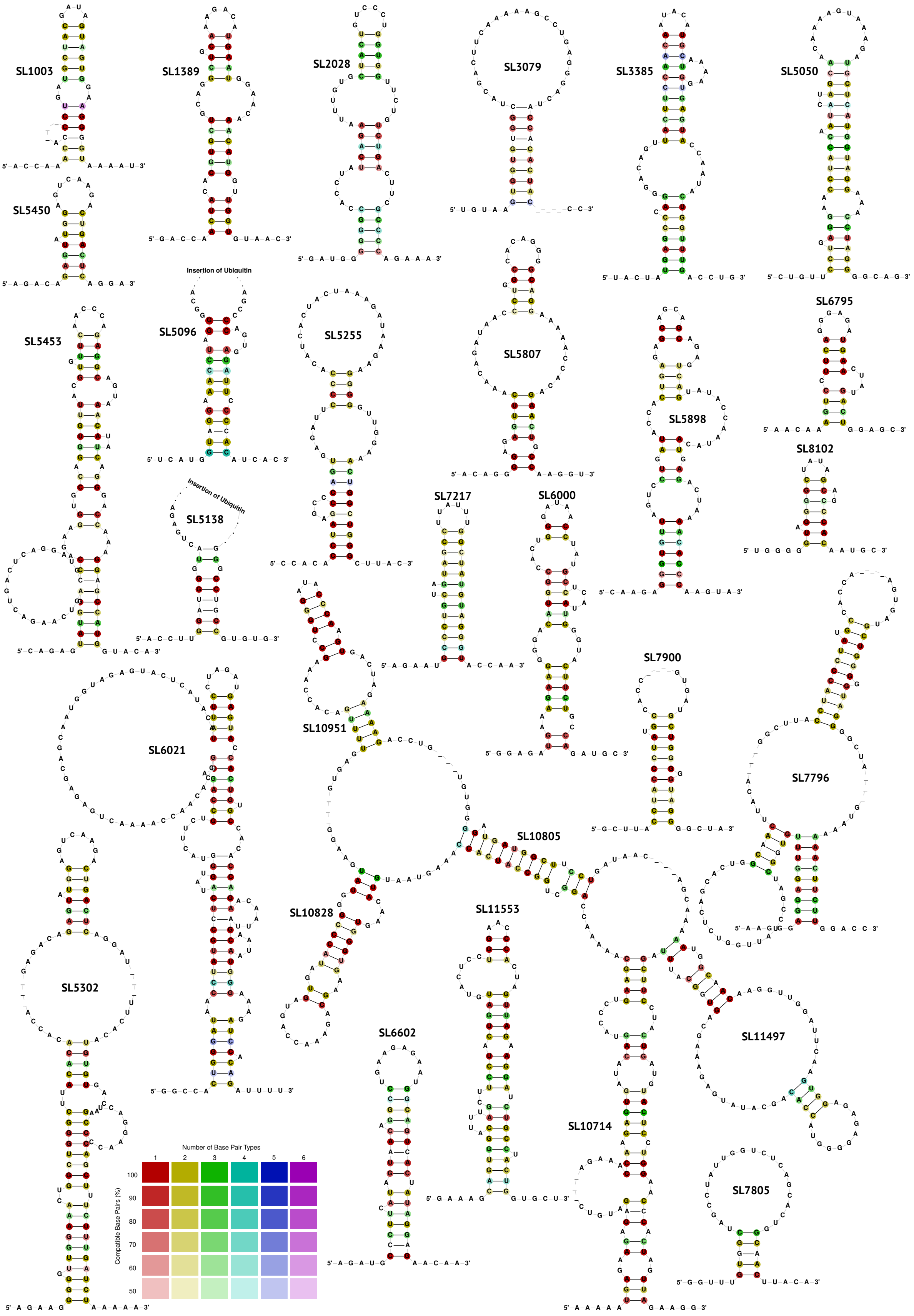


**Figure S7:** 29 novel conserved RNA secondary structure candidates from protein-coding region of the *Pestivirus* alignment.

### RNA secondary structures of species-subsets

**Table S3:** Conserved RNA secondary structures (SS) in a subset of the *Pestivirus* genus. Start and end positions refer to the alignment and reference strain for the corresponding species. We confirmed the previously predicted RNA secondary structures of SL II, SL III, and SL IV in the 3' UTR for species A and C separately (available in `Rfam` v15.1 or later); and predicted further 3 novel conserved RNA families (bold font) which will be incorporated into `Rfam`. We named novel structures according to their position in the reference genome.

| RNA SS | Genomic region | Alignment Position | | Genome Position | | Species (Reference Genome) |
|---|---|---|---|---|---|---|
| | | from | to | from | to | |
| **SL 637** | $N^{pro}$ | 1,186 | 1,224 | 637 | 675 | A (1-SD1, NC_076029), B, G, H, C, D, I, N, O, E, F, L, Wenzhou |
| **SL 4652** | NS2 | 5,675 | 5,728 | 4,652 | 4,695 | A (1-SD1, NC_076029), B, G, H, C, D, I, N, O, E, F, L, M, Wenzhou |
| **SL 5222** | NS3 | 6,863 | 6,889 | 5,222 | 5,248 | A (1-SD1, NC_076029), B, G, H, C, D, I, N, O, E, F, L, M, Wenzhou, P, Q, J, R, Jingmen |
| SL IV (RF04323) | 3' UTR | 13,947 | 14,142 | 12,114 | 12,129 | A (1-SD1, NC_076029) |
| SL II (RF04323) | 3' UTR | 14,194 | 14,693 | 12,141 | 12,233 | A (1-SD1, NC_076029) |
| SL IV (RF04324) | 3' UTR | 13,948 | 14,117 | 12,073 | 12,098 | C (Alfort/187, NC_038912) |
| SL III (RF04324) | 3' UTR | 14,119 | 14,240 | 12,100 | 12,174 | C (Alfort/187, NC_038912) |
| SL II (RF04324) | 3' UTR | 14,365 | 14,694 | 12,175 | 12,227 | C (Alfort/187, NC_038912) |
| SL III (RF04325) | 3' UTR | 14,117 | 14,245 | 11,106 | 11,164 | K (NC_038964), S, Zikole |

### Potential Primer Sites

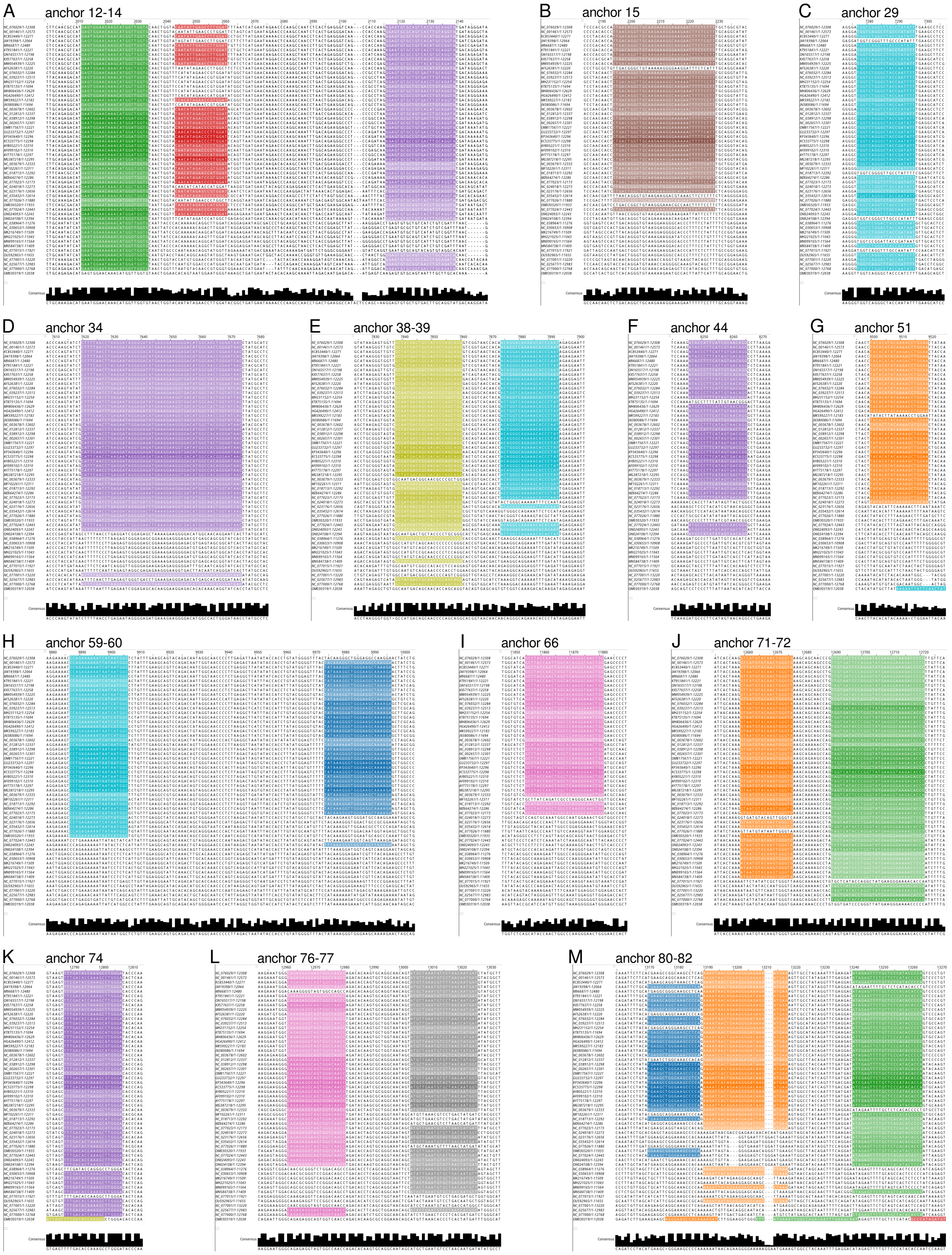


**Figure S8:** Potential primer sites for pestivirus detection identified by `AnchoRNA`.
